# Chromatin context-dependent regulation and epigenetic manipulation of prime editing

**DOI:** 10.1101/2023.04.12.536587

**Authors:** Xiaoyi Li, Wei Chen, Beth K. Martin, Diego Calderon, Choli Lee, Junhong Choi, Florence M. Chardon, Troy McDiarmid, Haedong Kim, Jean-Benoît Lalanne, Jenny F. Nathans, Jay Shendure

**Author notes:** Correspondence (X.L), (J.S.).

## Abstract

Prime editing is a powerful means of introducing precise changes to specific locations in mammalian genomes. However, the widely varying efficiency of prime editing across target sites of interest has limited its adoption in the context of both basic research and clinical settings. Here, we set out to exhaustively characterize the impact of the *cis-*chromatin environment on prime editing efficiency. Using a newly developed and highly sensitive method for mapping the genomic locations of a randomly integrated “sensor”, we identify specific epigenetic features that strongly correlate with the highly variable efficiency of prime editing across different genomic locations. Next, to assess the interaction of *trans*-acting factors with the *cis*-chromatin environment, we develop and apply a pooled genetic screening approach with which the impact of knocking down various DNA repair factors on prime editing efficiency can be stratified by *cis*-chromatin context. Finally, we demonstrate that we can dramatically modulate the efficiency of prime editing through epigenome editing, *i.e.* altering chromatin state in a locus-specific manner in order to increase or decrease the efficiency of prime editing at a target site. Looking forward, we envision that the insights and tools described here will broaden the range of both basic research and therapeutic contexts in which prime editing is useful.

## INTRODUCTION

Prime editing^1^ facilitates the precise installation of genetic variants with minimal off-target effects, including edits that are challenging to introduce with earlier generations of genome editing technologies^2–4^. The prime editor (PE), consisting of a fusion of a Cas9 nickase and a reverse transcriptase (RTase), searches for target site(s) and introduces a desired mutation with a prime editing guide RNA (pegRNA), which programs both the site specificity and the nature of the mutation. The high programmability and specificity of prime editing, together with its avoidance of double-stranded breaks, offers considerable advantages over alternatives in the context of therapeutic genome editing^5^. Its programmability and specificity have also facilitated other goals, such as sequential genome editing for molecular recording^6–8^ and the high-throughput functional characterization of genetic variants^9^.

A counterpoint to this promise is that the efficiency of prime editing at an endogenous locus is generally low and highly variable across target sites^1, 10–12^. Factors impacting this efficiency are likely to include: 1) the properties of the prime editing ribonucleoprotein (RNP) complex itself; 2) the primary sequence of the target and programmed edit; 3) *trans-*acting factors such as the endogenous DNA repair proteins involved in the installation of edits; and 4) the *cis*-chromatin context where the target resides. The first three of these factors have been explored to varying extents, with insights facilitating improvements to prime editing efficiency. For the first factor, an engineered RTase (PE2)^1^, a codon- and structure-optimized prime editor protein (PEmax)^11^, and a structurally stabilized pegRNA (epegRNA)^13^ have been developed. For the second factor, the impact of the target sequence and the edit sequence has been examined in the context of various kinds of edits (*e.g.* substitutions and indels), and corresponding machine learning models have been generated to facilitate design^10, 14, 15^. For the third factor, a rate-limiter for prime editing is DNA mismatch repair (MMR), which detects the intermediate product of prime editing and efficiently repairs the edited strand. Through mechanistic understanding or screens of repair-relevant *trans-*acting factors, strategies that enhance prime editing efficiency, either by misleading the MMR machinery by nicking the non-edited strand (PE3)^1^ or by transiently dampening the MMR pathway^11, 12^, have been reduced to practice.

In contrast, the role of the fourth factor, *cis*-chromatin environment, in modulating the efficiency of prime editing remains largely uncharacterized. Mammalian genomes are packaged into chromatin. Chromatin compaction is known to physically shield DNA from damage caused by DNA damaging agents^16^. Various epigenetic factors can interact with DNA repair factors to modulate the kinetics of DNA damage response (DDR)^17–19^. In the context of CRISPR-based gene editing, activity of Cas9 is strongly influenced by chromatin. Nucleosomes inhibit the binding and target cleavage of Cas9^20, 21^. The epigenetic environment not only influences the overall efficiency but also alters the balance of DNA repair pathways that are used to repair a double-strand break (DSB)^22–25^. For example, while non-homologous end joining (NHEJ) is the major DSB repair mechanism, targets in heterochromatin regions have a slightly higher preference for the microhomology-mediated end joining (MMEJ) pathway^23^. Prime editing, which is also a CRISPR-based gene editing method and involves the generation and repair of a single-strand DNA break, is likely to also be impacted by the chromatin state of the target sequence and its flanking regions.

In this study, we first sought to determine the impact of chromatin context on prime editing outcomes. We randomly integrated a large number of chromatin “sensors” carrying synthetic prime editing targets throughout the genome. To determine the genomic locations of these sensors, we developed and optimized a T7-assisted reporter mapping assay, which facilitates mapping of the precise locations of thousands of reporters with high sensitivity. Using this approach, we observe highly variable prime editing outcomes across genomic contexts, and identify epigenetic features that are predictive of both high and low prime editing efficiency. Importantly, when the target site falls within a gene, prime editing efficiency is positively correlated with the transcriptional activity of that gene, but negatively correlated with the distance between the target and transcription start site (TSS).

The observed variation in prime editing efficiency reflects the compound effect of differential chromatin accessibility for the prime editing complex and site-specific regulation of DDR. Therefore, we next sought to dissect chromatin context-dependent regulation of prime editing using functional genomic screens. Recently, several DDR gene-focused screens, including a high-throughput pooled genetic screen, have been carried out in the context of prime editing, pinpointing the key role of the MMR pathway in this process^11, 12^. However, neither of these studies were designed to characterize the role of the local chromatin environment in modulating cellular response to perturbation in the DDR pathways, *i.e.* the intersection of *trans-*acting DDR factors with the *cis-*chromatin environment. To get at this question, we developed a pooled genetic screen, based on a highly scalable method for single cell transcriptional profiling (sci-RNA-seq3)^26, 27^. By incorporating a T7 *in situ* transcription (IST) reaction, we repurposed the sci-RNA-seq3 method to profile prime editing outcomes in pre-integrated reporters across the genome. With this method, we profile the impact of a library of shRNAs against 74 DDR-related genes on 50 synthetic prime editing reporters with defined locations, and identify previously uncharacterized factors that play chromatin context-dependent roles in regulating prime editing outcomes.

Finally, since prime editing outcomes are highly influenced by the local chromatin environment, we explored the possibility of modulating prime editing efficiency by first editing the epigenome of the target region^28, 29^. Leveraging CRISPRoff, a recently developed epigenetic reprogramming tool^28^, we find that prime editing efficiencies in intragenic targets were attenuated after epigenetic silencing of the gene in which they reside, further corroborating the link between transcriptional activity and prime editing outcomes. Building on this observation, we also show that transient gene activation by CRISPRa before prime editing of endogenous genomic sequences can dramatically enhance editing efficiency, demonstrating a new avenue for enhancing prime editing outcomes with potential therapeutic utility.

## RESULTS

### Design and efficient mapping of prime editing reporters by T7 *in vitro* transcription

To enable the measurement of *cis*-chromatin effects on prime editing efficiency, we developed a strategy for mapping the precise genomic locations of densely integrated reporters across the genome (**Fig. 1A**). The method relies on linear amplification of insertion junctions by *in vitro* transcription (IVT) from an internal T7 promoter in purified genomic DNA (gDNA). After gDNA removal, *in vitro* transcribed RNAs were purified and reverse transcribed using RT primers with 8-bp degenerate ends. Then, cDNAs were amplified by semi-specific PCR and sequenced by paired-end sequencing to associate construct barcodes with their neighboring genomic sequences. Due to random priming of RT primers, only sequencing reads spanning insertion junctions were used for inferring precise integration sites at single-base resolution (**Supplementary Fig. 1A**). Compared to conventional reporter mapping approaches (*e.g.* inverse-PCR^30^, Splinkerette PCR^31^, LAM-PCR^32^ and tagmentation-assisted PCR^33^), this method has two key advantages. First, it does not rely on biochemical steps that are biased and display suboptimal efficiency (*e.g.* digestion and ligation). Second, the highly processive T7 polymerase amplifies molecules carrying positional information before capture with RT primers, which further increases sensitivity.

**Figure 1.**
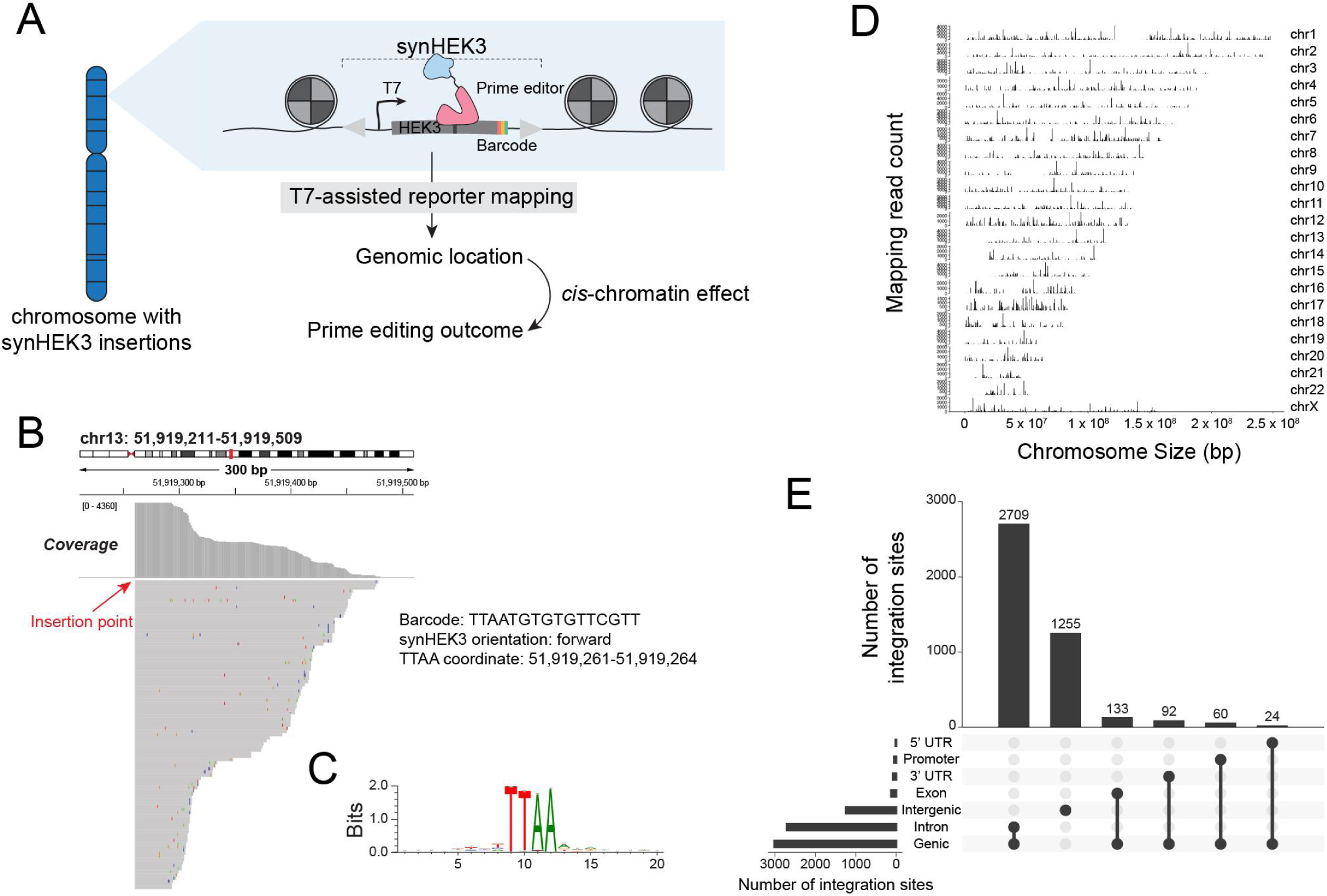
Efficient genome-wide mapping of the precise integration sites of a synHEK3 chromatin sensor. **A**) Measuring *cis*-chromatin effects on prime editing efficiency. A library of prime editing reporters (synHEK3) is randomly inserted across the genome. Genomic locations of these reporters are determined by a T7-assisted reporter mapping method. The *cis*-chromatin contexts of the mapped reporters are used to model prime editing outcomes as measured from each reporter. **B**) Genome browser view of a read pileup pinpointing the location to which a synHEK3 chromatin sensor is inserted in chromosome 13. The inferred insertion junction is marked. Barcode, orientation and coordinates of the synHEK3 reporter are annotated. **C**) Motif enrichment analysis of 20-bp windows surrounding synHEK3 insertion sites. **D**) Coverage plot of the unique synHEK3 reporters identified in the 500-cell pool (n = 4,273). **E**) Genomic annotation and counts of synHEK3 reporters (counting only the subset marked by unique barcodes).

We developed a compact *piggyBac*-based chromatin “sensor”, which contains a synthetic bacteriophage T7 promoter, the target sequence for a highly efficient spCas9 guide RNA (gRNA) – *HEK3*, and a 16-bp barcode (BC) serving as a unique molecular identifier for the reporter and its genomic insertion site. The final construct was 358 base pairs in size including two short inverted terminal repeats (ITRs) and lacks any known regulatory elements that could potentially interfere with the local chromatin environment (**Supplementary Fig. 1A**). We randomly integrated a complex library of the synthetic PB-T7-HEK3-BC (designated synthetic HEK3 or synHEK3) reporters into a monoclonal K562 cell line constitutively expressing the PE2 prime editor^2^ (**Supplementary Fig. 1B**). After the *piggyBac* transposase was diluted out and all synHEK3 integrations were stable, we measured synHEK3 reporter copy number by qPCR. On average, 15.5 copies of the synHEK3 reporters were detected per cell (**Supplementary Fig. 1C**). We further bottlenecked the pool to ∼500 clones, each containing a unique combination of randomly inserted reporters, and performed downstream measurements on cells derived from this population (**Supplementary Fig. 1B**).

We first sought to determine the genomic locations of the integrated synHEK3 reporters in the bottlenecked K562 pool using the T7-assisted reporter mapping method. Aligned reads formed clusters with sharp boundaries when visualized on a genome browser, such that the boundary corresponds to the putative insertion point of the chromatin sensor at base-pair resolution (**Fig. 1B**). After read counting and barcode error correction (**Methods**), we identified 10,095 insertion sites (**Supplementary Fig. 1D**). The mapping was near-complete, as supported by saturation analysis upon over-sequencing (**Supplementary Fig. 1E**). Motif analysis of sequences around the insertion sites revealed a highly specific TTAA motif at the insertion junction (**Fig. 1C**), consistent with the expected insertion motif for the *piggyBac* transposon^34^. We removed sites that did not intersect with a TTAA motif (N = 645/10,095, 6.4%) and used the genomic coordinates of the TTAA motifs as locations of the synHEK3 reporters. Of note, despite the large number of insertion sites identified in this pool, only 4,273 (42.3%) sites were marked by unique barcodes (**Fig. 1D**), and 880 barcodes were found associated with more than one integration site, indicating frequent *piggyBac* transposon excision and re-integration events after DNA replication but before the *piggyBac* transposase was fully diluted out (**Supplementary Fig. 1F**). Annotation of these sites suggests *piggyBac* integration is biased towards accessible regions such as promoters and 5’ UTRs (**Fig. 1E**; **Supplementary Fig. 1G,H**).

### Impact of chromatin features on prime editing efficiency

With the K562 reporter pool established and characterized, we next sought to measure prime editing outcomes across different genomic contexts. We transiently transfected the reporter cell population with a pegRNA designed to install a CTT trinucleotide insertion at the *HEK3* loci^1^. After 4 days, we performed amplicon sequencing of the synHEK3 sites to estimate editing efficiencies for all uniquely barcoded reporters (**Fig. 2A**). This pegRNA empirically drove a 17% editing rate at the endogenous *HEK3* loci. However, a very broad range of editing efficiencies (0-94%) were observed when the same target sequence was integrated at different genomic locations, suggesting prime editing outcomes are highly influenced by the local chromatin environment (**Fig. 2A**).

**Figure 2.**
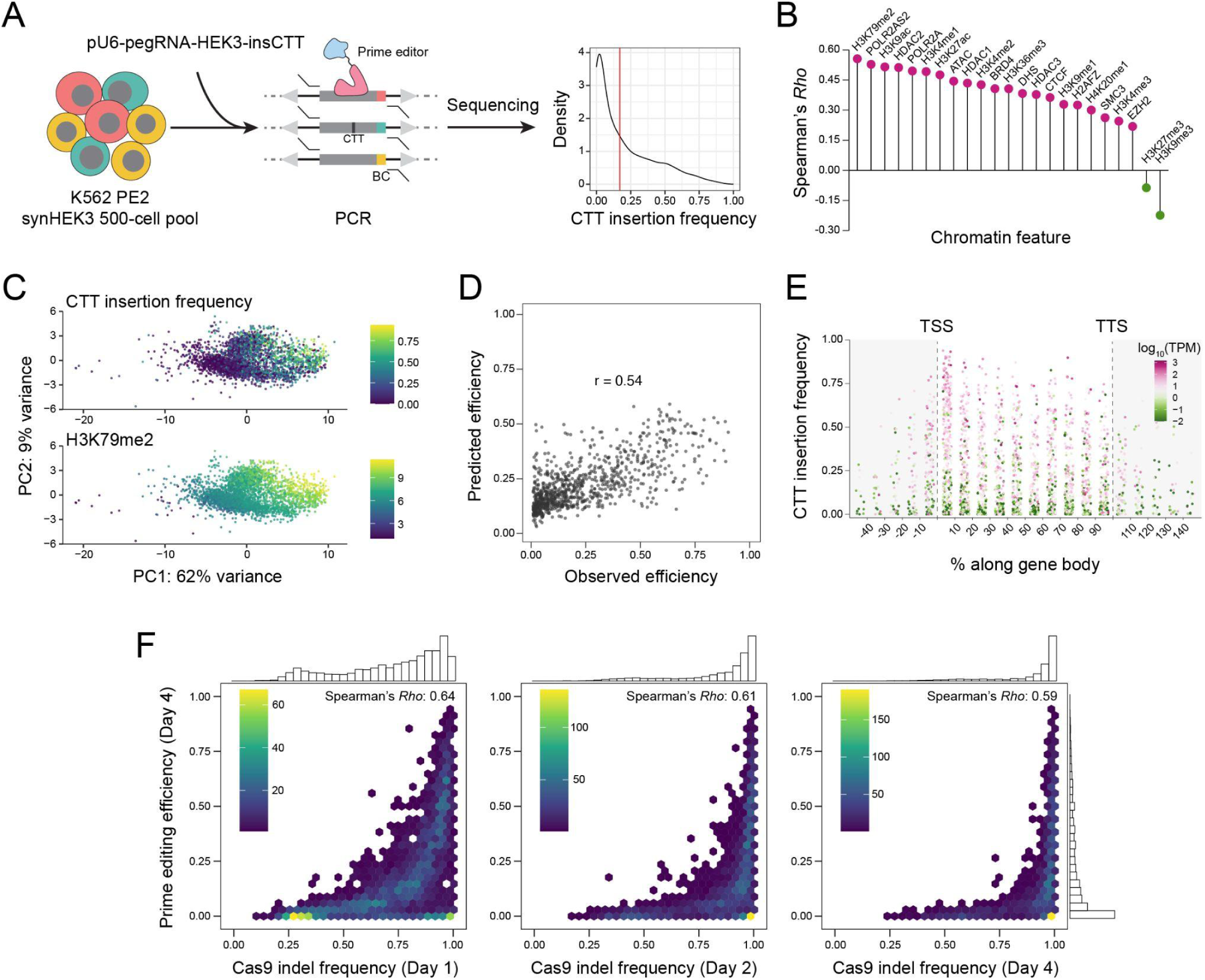
Chromatin context has a major impact on prime editing efficiency. **A**) Left: the K562 PE2 synHEK3 500-cell pool was transfected with the pU6-pegRNA-HEK3-insCTT plasmid. After 4 days, prime editing efficiency was measured by amplicon sequencing of synHEK3 reporters. Right: density plot of CTT insertion frequency in all uniquely barcoded synHEK3 reporters. Red line indicates empirical editing efficiency (17%) at the endogenous *HEK3* loci in the same cell line. **B**) Spearman correlation (*Rho*) between prime editing efficiencies and chromatin feature scores of the 2-kb window surrounding each reporter’s integration site. **C**) PCA plots of synHEK3 reporters generated using ENCODE-derived chromatin scores as features. The first two principal components (PCs) are plotted (PC1: variance 62%; PC2: variance 9%). Points are colored by prime editing efficiency (top) or the ChIP-seq signal of H3K79me2 (bottom). **D**) Scatter plot of observed and predicted prime editing efficiencies by the beta regression model using reporters in a holdout test set. **E**) Prime editing efficiency for gene-proximal reporters. Distance was calculated relative to the closest TSS, scaled by gene length and binned. Points are colored based on expression levels (log_10_) of the genes. TPM, transcripts per million. **F**) Comparison of total editing efficiency for Cas9 vs. prime editing, using the same pool of synHEK3 reporters at Days 1, 2 and 4 after Cas9 RNP transfection. Plots are colored based on the density of reporters underneath. 1-D histograms of total editing efficiencies are plotted at the top and right sides.

To assess which chromatin features are associated with the enhancement or suppression of prime editing, we inspected a series of epigenetic datasets generated from K562 cells by the ENCODE Consortium^35^, including DNase hypersensitive sites (DHS), ATAC-seq, and ChIP-seq datasets for all available histone modifications, as well as selected DNA binding proteins and epigenetic modulators (**Table S2**). We focused on 2-kb windows centering on the sites in which chromatin sensors were integrated, and computed scores over these windows for the selected epigenetic datasets (**Methods**; **Table S3**). Prime editing efficiency broadly correlated with markers of euchromatin. Among these, H3K79me2 levels were most highly correlated with prime editing efficiency (Spearman’s *Rho*=0.56; **Fig. 2B-C**; **Supplementary Fig. 2A**). Finally, we trained a beta regression model using these epigenetic scores. On a holdout test set, the model predicted prime editing efficiencies with a pseudo r of 0.54 (**Fig. 2D**). In this model, H3K79me2 levels had the largest positive coefficient, which again suggests that at least among the available epigenetic data, it is the strongest predictor of prime editing efficiency.

In mammalian cells, H3K79 methylation is solely deposited by DOT1L, which is found in proximal regions of active genes and coupled with active transcription^36–38^. This is consistent with the fact that H3K79me2 is primarily observed in early transcribed regions^37^ and its ChIP-seq signal is more spread out than H3K4me3, a marker of promoters (**Supplementary Fig. 2B**). Therefore, we hypothesized that the prime editor generally favors transcriptionally active regions and is most proficient immediately downstream of TSSs. To evaluate this further, we computed distances of the chromatin reporters to the nearest TSSs and their corresponding transcription termination sites (TTSs), and then binned them by proportional position along the gene body. We observe that chromatin reporters immediately downstream of promoters had the highest prime editing rates. The difference was more striking for synHEK3 reporters near highly expressed genes (**Fig. 2E**; **Supplementary Fig. 2C**).

### Implications of Cas9 editing outcomes at the same synHEK3 reporters

The observed wide range of prime editing efficiencies at synHEK3 reporters throughout the genome could be due to differential accessibility for the prime editing complex. We wanted to determine whether the binding of Cas9 alone was similarly affected. We transfected the K562 synHEK3 reporter pool with Cas9 RNP bearing a gRNA targeting the synHEK3 reporters (*i.e.* the same spacer sequence as the synHEK3-targeting pegRNA described above), and compared editing outcomes between prime editing and Cas9 editing. This experiment confirmed that Cas9 editing is much more efficient than prime editing. Frequent indels were observed as early as Day 1, and 1,886 out of 2,517 (75%) reporters had indel frequencies higher than 90% at Day 4. Furthermore, Cas9 was able to mutate synHEK3 reporters that were relatively “resistant” to prime editing (**Fig. 2F**; **Supplementary Fig. 3A**). As expected, synHEK3 reporters near highly expressed genes were more frequently edited by Cas9. When inspecting the activity of Cas9 near highly expressed genes, we found that reporters immediately downstream of TSSs had higher indel frequencies at Day 1 (similar to what was observed for prime editing at Day 4; **Fig. 2E**), but the differences were negligible at later time points. On Day 4, reporters in gene-proximal regions were edited to near-saturation by Cas9 (**Supplementary Fig. 3B**). The most immediate response to Cas9 RNP at Day 1 reflected the impact of chromatin accessibility on Cas9 binding, which might similarly shape the binding of the prime editing complex. This suggests that the H3K79me2 mark might define a region (more specific than the more broadly defined accessible regions revealed by DNase hypersensitivity or ATAC-seq) that is immediately downstream of the TSS of expressed genes and allows for efficient binding of Cas9 or the prime editor.

Repair of Cas9-induced DSBs also results in diverse outcomes, providing us with rich information for dissecting the impact of the chromatin landscape on DNA repair kinetics. Previously, this relationship was explored using the “thousands of reporters integrated in parallel” (TRIP) strategy^39^. Cas9-induced DSBs are mainly repaired through the NHEJ and MMEJ pathways, and MMEJ is known to have slower repair kinetics compared to NHEJ^40^. Increased preference for MMEJ for repairing a DSB was observed for reporters residing in heterochromatic regions^23^. We computed the balance between MMEJ and NHEJ as MMEJ / (MMEJ + NHEJ) ratios^23^ for the synHEK3 reporters at Day 4 and corroborated this observation (**Supplementary Fig. 3C**). Moreover, we found that even within genes, the MMEJ / (MMEJ + NHEJ) ratio steadily increased towards the TTS (**Supplementary Fig. 3D**), which reflected the concomitant increase of MMEJ allele frequencies and decrease of NHEJ allele frequencies (**Supplementary Fig. 3E**). The differential preference for the two pathways along the gene body suggests a gradient effect posed on DNA break repair kinetics by chromatin such that faster repair mechanisms are favored immediately downstream of TSSs. A similar TSS-distance gradient may affect which repair mechanisms ultimately resolve prime editing-induced single strand breaks (SSBs).

### Strategy for investigating the chromatin context-dependencies of *trans-*acting factors influencing prime editing

In addition to Cas9 binding, DNA damage repair is tightly regulated by local chromatin context^17^. To explore potential mechanisms mediating chromatin context-specific regulation of prime editing, we decided to perform a DDR-focused genetic screen on synHEK3 reporters inserted at different genomic contexts, via a Perturb-seq style approach^41–43^ in which perturbations are coupled to outcomes via single cell molecular profiling. To co-profile the genetic perturbation(s) received by a given cell and the prime editing outcome recorded at its synHEK3 reporters, we sought to use T7 IST^44, 45^ to drive RNA production from the synHEK3 reporters in fixed, permeabilized nuclei, such that we could then capture these RNA molecules via a modified version of a scalable method for single cell transcriptional profiling, sci-RNA-seq3^26^.

We focused our analysis on synHEK3 reporters found in two monoclonal K562 lines derived from the original pool (referred to as clone 3 and clone 5 hereafter). The numbers of uniquely barcoded reporters were 22 and 28, respectively, and these reporters spanned major epigenetic features characterized above (**Fig. 3A**; **Table S4**). Prime editing efficiencies measured in these monoclonal lines were highly correlated with those estimated for the same reporters in the original polyclonal population, indicating robustness of the chromatin sensors (**Supplementary Fig. 4A**).

**Figure 3.**
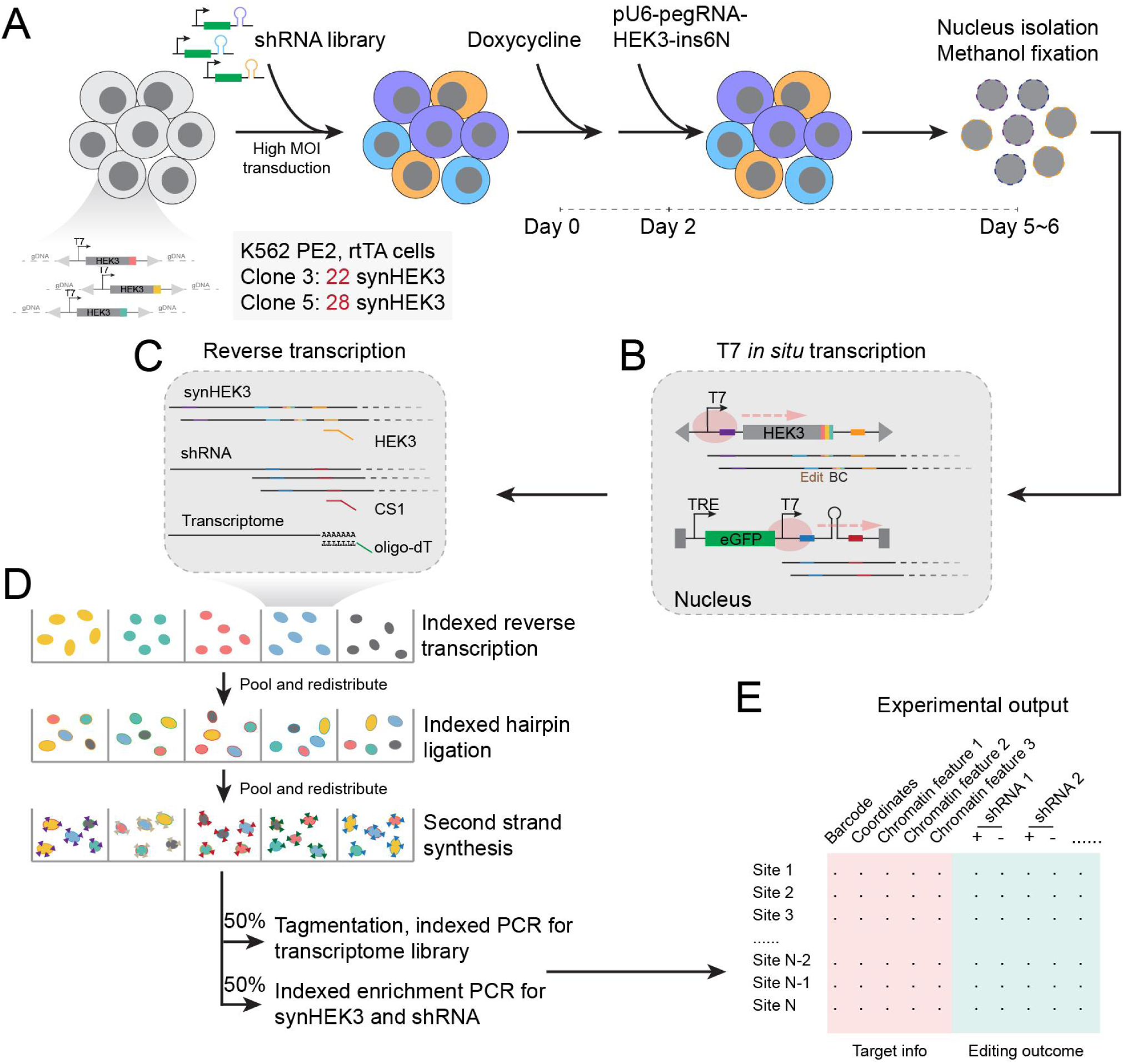
Dissecting chromatin context-dependent regulation of prime editing using a modified sci-RNA-seq3 workflow. Experimental workflow of the pooled shRNA screen. **A**) The two monoclonal K562 lines used in this experiment stably expressed PE2 and rtTA, and together contained 50 unique synHEK3 reporters. Cells were transduced with the shRNA library at a high MOI manner and treated with Doxycycline (1 ug/mL). On Day 2, cells were transfected with pegRNAs to induce random 6-bp insertions. After 3-4 days, nuclei were extracted and fixed with ice-cold methanol. **B**) Fixed nuclei were subjected to IST with T7 polymerase (pink circle) to produce transcripts from the synHEK3 and shRNA constructs. **C**) Nuclei were distributed to 96-well for indexed RT (the first index). In each well, a cocktail of three indexed RT primers were used: oligo-dT primers, and synHEK3- and shRNA-specific primers. **D**) After RT, nuclei were pooled and redistributed into 96-well plates for indexed hairpin ligation (the second index). Then, they were pooled and split to final 96-well plates. After second-strand synthesis, nuclei were lysed and the resulting lysates were split to two plates. One plate underwent Tn5 tagmentation and indexed PCR to generate a transcriptome library. The other plate was used for indexed enrichment PCR to amplify the synHEK3 and shRNA constructs. **E**) Expected output of the experiment. For every synHEK3 reporter, editing outcomes were computed and compared between cells with vs. without a specific shRNA.

For genetic perturbation, we chose an miR-E based shRNA system^46^, which is able to trigger a potent knockdown effect at a single copy and is orthogonal to prime editing. In this system, an shRNA is embedded in an optimized miRNA backbone within the 3’ untranslated region (3’ UTR) of a GFP expression cassette driven by a tetracycline-controlled promoter^46, 47^. We constructed our shRNA library by selecting 4 shRNAs for each candidate gene (**Table S5**)^47^. The library contained 304 shRNAs against 76 genes, including 74 DDR-related genes (10 without detectable expression) and 2 luciferase genes. shRNAs against unexpressed genes and luciferase genes served as internal controls for this experiment. The 64 DDR-related, expressed genes comprised hits found by Repair-seq^11, 48^, genes involved in other major DNA damage repair pathways, and epigenetic factors involved in H3K79me2 metabolism (**Supplementary Fig. 4B**). To enable post-fixation enrichment and targeted capture of the shRNA transcripts in sci-RNA-seq3, we further modified the lentiviral construct to contain a T7 promoter upstream of the shRNA and a RT primer binding site (PBS) (**Supplementary Fig. 4C**). Before the experiment, we engineered the two monoclonal lines to stably express a reverse tetracycline-controlled transactivator (rtTA). We then transduced the cells with the lentiviral library at a multiplicity of infection (MOI) of 10 and removed any non-transduced cells by antibiotic selection (**Fig. 3A**).

We treated the cells with Doxycycline for 2 days to fully induce knockdown effects. We then transfected pegRNAs encoding random 6-bp insertions. The use of random insertions allowed us to minimize potential bias caused by the insertion sequence itself^49^, as well as to distinguish real editing outcomes (*in situ* transcribed in nuclei) from ambient transcripts. After 3-4 days, perturbed cells derived from the two clones were mixed at a 1:1 ratio and profiled leveraging a modified sci-RNA-seq3 workflow (**Fig. 3A**). The first modification to the protocol occurred before the RT step. We performed T7 IST on methanol-fixed nuclei to drive expression of synHEK3 and increase transcript abundance of the shRNAs (**Fig. 3B**). Second, at the RT step, a cocktail of an oligo-dT primer and primers specific for the synHEK3 and shRNA transcripts were used to simultaneously capture the cellular transcriptome, editing outcome, and perturbation status (**Fig. 3C**). After the third round of nucleus splitting and second strand synthesis, nuclei were lysed, with one half of the lysate undergoing normal sci-RNA-seq3 transcriptome library preparation and the other half of the lysate subjected to enrichment PCR for both the synHEK3 and shRNA constructs (**Fig. 3D**; **Methods**).

For each experiment, three libraries were sequenced: a sci-RNA-seq3 library of the cellular transcriptome, an shRNA library, and a synHEK3 library bearing reporter barcodes and prime editing outcomes (**Supplementary Fig. 4D**). The transcriptome library was only shallowly sequenced and used to determine high-quality cells. The captured shRNA and synHEK3 transcripts were assigned to these cells based on the matching combinatorial indices. From this, we generated an shRNA recovery matrix for all cells. For every cell-synHEK3 reporter pair, we determined its editing outcome based on its sequence change and generated an editing outcome matrix for all such pairs. We then applied a quantitative trait locus (QTL)-like analysis to the data. More specifically, for each synHEK3-shRNA pair, we aggregated cells receiving the corresponding shRNA and compared the editing frequency of this synHEK3 reporter against that of the same synHEK3 reporter in cells without this shRNA (**Fig. 3E**).

### Identification of *trans*-acting regulatory factors of prime editing

We focused our analysis on a set of high-quality cells (n = 667,810) as assessed from the sci-RNA-seq3 transcriptome library (**Methods**). Reads derived from synHEK3 reporters from the two clones were well-separated (**Supplementary Fig. 5A**). After a series of filtering steps (**Methods**), we retrieved a total of 229,657 cells (105,583 and 124,074 for clones 3 and 5, respectively), for which at least one shRNA and one synHEK3 construct were detected. The median numbers of cells per shRNA in clone 3 and clone 5 were 1,604 and 2,208, respectively, and the median numbers of shRNAs detected per cell were 4 and 5, respectively (**Supplementary Fig. 5B**). For synHEK3, the median numbers of barcodes detected per cell were 7 and 11, respectively (**Supplementary Fig. 5B**). We next collapsed reads on a per-cell, per-barcode basis such that we had a single read representing each individual cell for downstream editing efficiency analyses of the synHEK3 reporters. To determine whether we could reliably estimate editing efficiency using the collapsed reads from single cells, we sampled a small fraction of the cells in this screen and performed amplicon sequencing of the synHEK3 reporters in parallel. Prime editing efficiencies estimated by aggregating single cells were highly accurate and almost perfectly correlated with those estimated by bulk amplicon sequencing (**Supplementary Fig. 5C**).

To assess whether the experimental setup empowered us to detect shRNA-mediated knockdown effects on prime editing outcomes, we calculated and compared editing efficiency in cells with a given candidate shRNA vs. cells without that shRNA (control cells), repeating this for every synHEK3-shRNA combination. To statistically assess this, we treated prime editing outcomes as binary events (unedited vs. insertion) and performed a binomial test in cells with a given shRNA. Using editing efficiency estimated in control cells (*i.e.* all cells without that shRNA) as the null hypothesis, we computed a statistical significance value (*p* value) for the observed editing outcome. We iteratively performed binomial tests for all synHEK3-shRNA combinations in the two monoclonal lines and observed a clear excess of significant synHEK3-shRNA pairs compared to pairs involving control shRNAs (**Fig. 4A**). 337/7,168 and 250/5,632 synHEK3-shRNA pairs were significant at a false discovery rate (FDR) of 5% in clone 5 and clone 3, respectively. This excess of highly significant *p* values does not appear to be driven by differences in the number of cells associated with candidate-targeting shRNAs vs. control shRNAs (**Supplementary Fig. 5D**).

**Figure 4.**
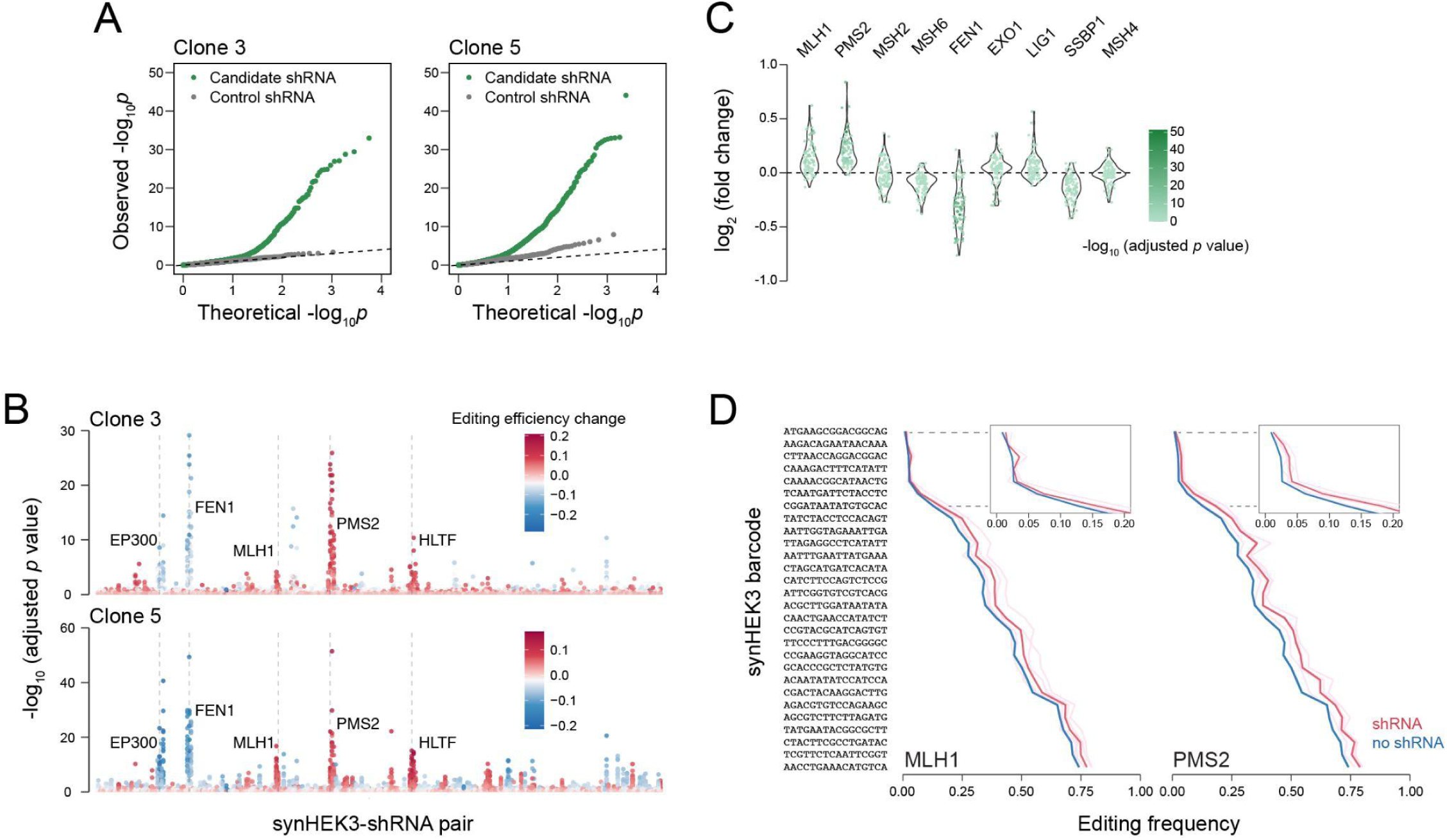
Effects of MMR pathway on prime editing. **A**) Q-Q plot of statistical significance (-log_10_) of synHEK3-shRNA pairs in clone 3 (left) and clone 5 (right). Candidate shRNAs (green) and control shRNAs (gray) are plotted separately. **B**) Plots of adjusted *p* values (-log_10_) of all synHEK3-shRNA pairs. Target genes with high statistical significance are annotated. Points are colored by editing efficiency changes caused by corresponding shRNAs. **C**) Effects of shRNAs targeting genes in the MMR pathway in clone 5. Log_2_ fold changes of prime editing efficiencies of synHEK3-shRNA pairs are plotted and colored by their corresponding adjusted *p* values (-log_10_). **D**) Effects of shRNAs against MLH1 and PMS2. Pink lines: editing frequencies in cells with individual shRNAs; red line: mean editing frequencies of the gene-targeting shRNAs; light blue lines: control editing frequencies for individual shRNAs (not visible because low variance relative to mean line, shown in blue); blue line: mean control editing frequencies.

Ordering *p* values by target gene identity revealed that shRNAs targeting several genes were highly significant, and reproducible across the two clones (**Fig. 4B**). The most prominent targets (*i.e.* those with multiple shRNAs passing the FDR threshold) include major components of the MMR pathway – PMS2, MLH1, and FEN1 – which have been previously reported to influence the installation of prime editing edits. To better visualize the effects of MMR proteins, we consolidated synHEK3-shRNA pairs involving shRNAs targeting MMR members included in this screen (**Fig. 4C**; **Supplementary Fig. 5E**). The strongest prime editing-promoting effects were observed for shRNAs against PMS2 and MLH1 (homologs of bacterial MutLα, **Fig. 4D**), which form a heterodimer and coordinate multiple repair steps after mismatch recognition^50^. Knocking down FEN1, a 5’ DNA flap endonuclease, led to strong suppression of prime editing in almost all synHEK3 reporters (**Supplementary Fig. 5F**), which is consistent with its reported role^11^. Perturbing the bacterial MutSα homologs^50^ MSH2 and MSH6 only yielded weak effects. MSH2 and MSH6 dimerize and preferentially recognize single base mismatches and 1-2-bp insertions and deletions. In clone 5, downregulation of MSH6 even led to a slight decrease in prime editing efficiency. Besides MSH6, MSH2 also dimerizes MSH3 and detects long insertions or deletions (>2 bp). Knocking down MSH6 potentially released the sequestered MSH2 and allowed for more efficient detection and correction of the 6-bp insertions introduced in this experiment. In contrast to the reported roles of EXO1 and LIG1 in prime editing-mediated single-nucleotide substitutions^11, 12^, perturbing these two genes did not alter editing efficiency of 6-bp insertions. Moreover, knocking down the single-strand DNA binding protein SSBP1 caused a slight decrease in prime editing efficiency in clone 5. Finally, knocking down another MutS homolog MSH4, which is involved in meiotic recombination^50^ and is not expressed in K562 cells, didn’t cause significant changes across synHEK3 reporters in either clone (**Fig. 4C**; **Supplementary Fig. 5E**).

In addition to MMR genes, we found that downregulation of EP300 negatively impacted prime editing efficiencies across all sites (**Supplementary Fig. 5G**). EP300 encodes the p300 histone acetyltransferase protein that works as an important transcriptional coactivator^51^. Its direct involvement in DNA damage response remains elusive. Furthermore, we cannot rule out the possibility that the observed effects of targeting EP300 result from downregulation of global transcriptional output including expression of the prime editor complex.

### HLTF works as a chromatin context-dependent repressor of prime editing

Another strong hit identified in the screen was HLTF, which is a member of the SWI/SNF2 family and contains an E3 ubiquitin ligase domain. HLTF mediates polyubiquitination of PCNA, which promotes replication through DNA lesions in an error-free manner^52^. A previous report has shown that knocking down HLTF led to a slight decrease in the frequency of prime editing-mediated single-nucleotide substitutions^11^. In contrast, we observed strong upregulation of 6-bp insertion frequencies, which was comparable to editing frequency changes induced by shRNAs against MLH1 and PMS2, suggesting potential differences in mechanisms involving HLTF in the repair of short and longer mismatches (**Fig. 5A**).

**Figure 5.**
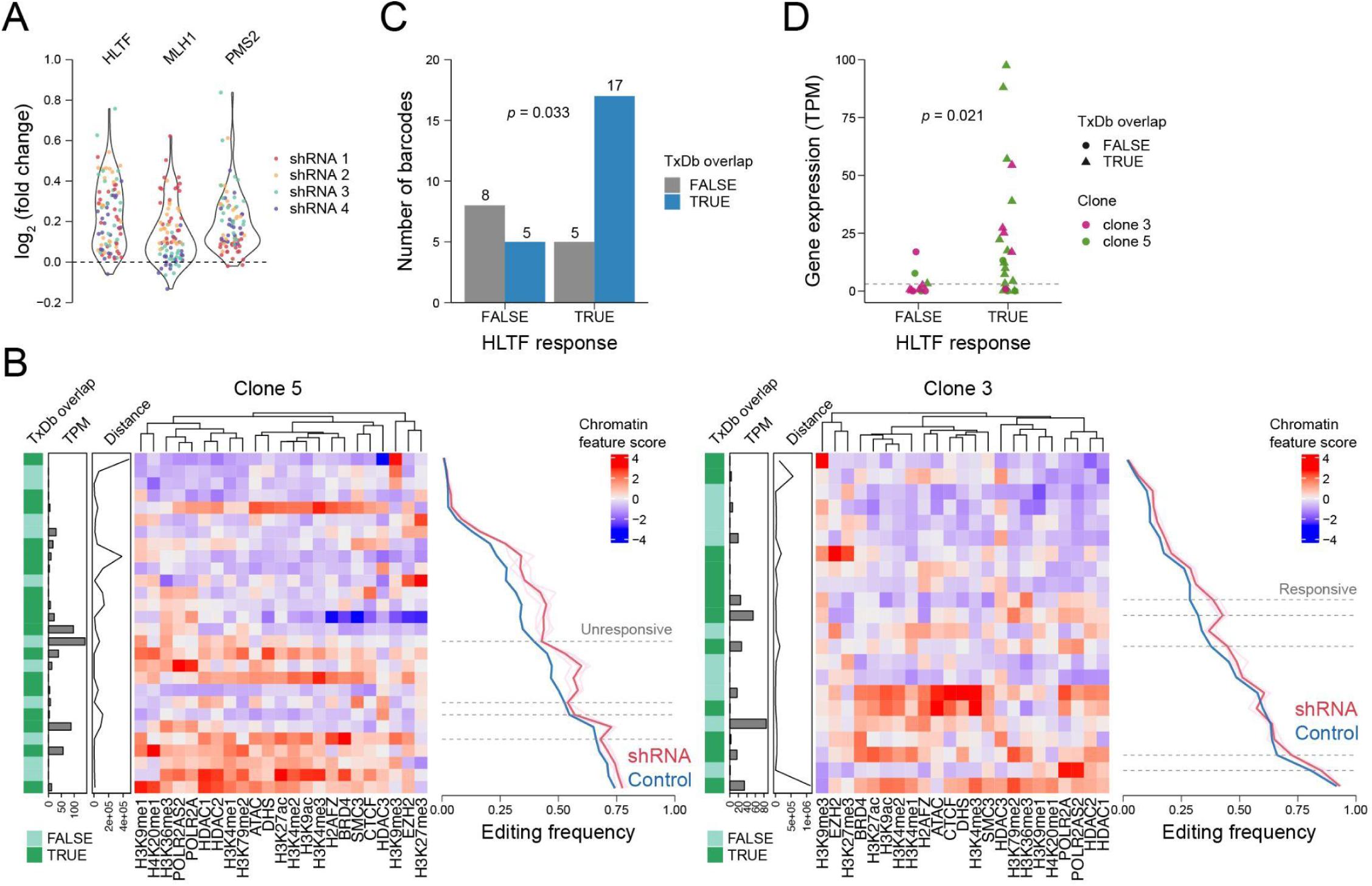
Chromatin context-specific response to HLTF inhibition. **A**) Violin plot of synHEK3 responses to inhibition of HLTF, MLH1 and PMS2. Points are colored by shRNA identity. **B**) Heatmap of synHEK3 reporters (row) and their responses to shRNAs against HLTF (left: clone 5; right: clone 3). The leftmost bar annotates the overlapping status of synHEK3 reporters with the GRCh38 TxDb records. The second left bar indicates the expression status of the overlapping gene or nearest gene in TPM. The third left bar indicates distances (in bp) to corresponding TSS of gene-overlapping synHEK3 reporters or distances to nearest genes for synHEK3 reporters outside genes. The middle heatmap is generated using scaled epigenetic scores of the synHEK3 reporters and clustered by column. The line plot on the right shows effects of shRNAs against HLTF. Pink lines: editing frequencies in cells with individual shRNAs; red line: mean editing frequencies of the gene-targeting shRNAs; light blue lines: control editing frequencies for individual shRNAs (not visible because low variance relative to mean line, shown in blue); blue line: mean control editing frequencies. Dashed lines: sites showing differential response to HLTF inhibition. TPM, transcripts per million. **C**) Bar plot of synHEK3 reporter counts based on their responsiveness to HLTF inhibition and overlapping status with the GRCh38 TxDb records. *P* value: Fisher’s exact test. **D**) Expression status (in TPM) of genes overlapping or proximal (within 10 kb) to synHEK3 reporter. Genes above the dashed line are considered expressed. *P* value: two-sided Kolmogorov–Smirnov test. TPM, transcripts per million.

Inspection of individual synHEK3 insertion sites revealed heterogeneous responses to HLTF inhibition (**Fig. 5B**; **Supplementary Fig. 6A**). Sites with low editing frequencies (< 0.2) showed little response. Due to editing frequencies being bounded on [0, 1], for sites with very high baseline editing frequencies, we likely underestimate the effects of the HLTF shRNAs due to the upper bound of the data. Therefore, for the following analysis, we only focused on synHEK3 reporters with baseline editing frequencies between 0.2 and 0.9. In clone 5, 4/21 of these sites showed much weaker increases in editing frequencies compared to reporters with similar starting editing frequencies. In clone 3, the synHEK3 reporters were overall less responsive to HLTF inhibition. However, 5/14 sites showed stronger and statistically significant upregulation of editing frequencies (**Supplementary Fig. 6A,C**). Of note, sites that were less responsive to HLTF inhibition showed slightly higher response to shRNAs against PMS2, and this effect was more obvious in clone 5 (**Supplementary Fig. 6B**). We grouped the selected synHEK3 reporters based on their responsiveness to HLTF shRNA knockdown (**Supplementary Fig. 6C**) and counted their overlapping status with annotated transcripts (genes). We found that the responsive group was enriched for sites overlapping with annotated genes (5.4-fold, fisher’s exact test *p* = 0.033; **Fig. 5C**). Furthermore, when taking expression status of the overlapping or nearby genes (within 10 kb) into consideration, the responsive sites tended to be found in the vicinity of actively transcribed genes, while the unresponsive sites were mostly found in non-transcribing regions (9.2-fold mean gene expression difference; *p* = 0.021; **Fig. 5D**).

To validate this result, we transduced clone 3 and clone 5 individually with constructs bearing either one of two shRNAs against HLTF or an shRNA against MLH1. After 2 days of doxycycline treatment to induce GFP and shRNA expression, we transfected the cells with pegRNAs encoding 6-bp insertions. We then compared log_2_ fold changes of editing frequencies between GFP-positive and negative cells. Because GFP (and presumably the shRNA embedded in its 3’ UTR) was heterogeneously induced, this effectively compared cells in which the knockdown is taking place vs. those in which it is not. The results confirm that the synHEK3 reporters showed differential responses to HLTF inhibition compared to MLH1 (**Supplementary Fig. 6D**).

In summary, by incorporating a T7 IST reaction into a Perturb-seq^41, 42^-like experiment in sci-RNA-seq3, we were able to associate genetic perturbations with phenotypes recorded in non-transcribing synHEK3 constructs, as well as to dissect the interaction between the *trans*-acting regulatory factors of prime editing and the *cis*-chromatin contexts. In addition to validating previously nominated modulators of prime editing and highlighting the mutation-dependencies of others, our data suggest that HLTF may work as a chromatin-context dependent repressor of prime editing.

### Modulating prime editing outcomes by targeted epigenetic reprogramming

The above experiments suggest a complex relationship between chromatin context and the efficiency of prime editing. Leveraging a constant “sensor” integrated to thousands of precisely mapped sites across the genome, we found that prime editing efficiency varies tremendously by genomic location (*e.g.* reproducibly 0% at some locations and 90% at others, despite the same edit being made to the same spacer-defined target sequence), with efficiency generally correlating with the open status of the chromatin (**Fig. 2B**). If the sensor was within a gene, its editing efficiency furthermore correlated with the transcriptional activity of that gene (**Fig. 2E**; **Supplementary Fig. 2C**). In addition, we pinpointed a histone mark, H3K79me2, that showed the highest correlation with prime editing efficiency (**Fig. 2B,C**), and characterized a negative correlation between the editing efficiency and distance to TSSs for intragenic targets (**Fig. 2E**; **Supplementary Fig. 2C**). Finally, through a functional genomics screen across many reporters, we showed that the locus-specific differences could be further impacted by site-specific regulation of DDR (**Figure 5**). Although locus-specific differences are not fully explainable by available annotations, these observations highlight the critical role of transcriptional activity in shaping how the epigenetic landscape modulates prime editing.

In a final set of experiments, we sought to ask whether or not we could exploit this relationship in order to alter the efficiency of prime editing, *i.e.* epigenetic reprogramming of the responsiveness of a target to prime editing. As a first step, we sought to examine whether the efficiency of editing of intragenic synHEK3 reporters could be altered by silencing the gene in which they resided. We chose three intronic synHEK3 reporters identified in the monoclonal line, clone 5, used above. These reporters were in the genes *USP7*, *METTL2A* and *LRRC8C*, and the distances between the reporters and corresponding TSSs were 3 kb, 6.5 kb and 1.1 kb, respectively (**Supplementary Fig. 7A**). For gene silencing, we used CRISPRoff, an epigenetic editing method that can establish highly specific long-term epigenetic repression of target genes^28^. In this experiment, we transiently transfected cells with plasmids encoding CRISPRoff-v2 and a pair of gRNAs targeting these genes’ promoters, with fluorescence-activated cell sorting used to enrich for transfected cells. On Day 11, we again transfected the cells with pegRNAs programming CTT insertions to all synHEK3 reporters (**Fig. 6A**). After calibrating control editing efficiency using the efficiencies of reporters that are not in target genes (**Fig. 6B**), we observed 39%, 26% and 47% decreases in editing frequencies of synHEK3 reporters in the genes *USP7*, *METTL2A* and *LRRC8C*, respectively (**Fig. 6C**). These reductions in prime editing efficiencies were likely direct effects of gene silencing as other synHEK3 reporters were not affected and RNA-seq indicated highly specific target gene suppressions (**Supplementary Fig. 7B**). Interestingly, the magnitude of editing efficiency reductions were on par with the magnitude of reductions in gene expression induced by CRISPRoff (53%, 30% and 55%, respectively).

**Figure 6.**
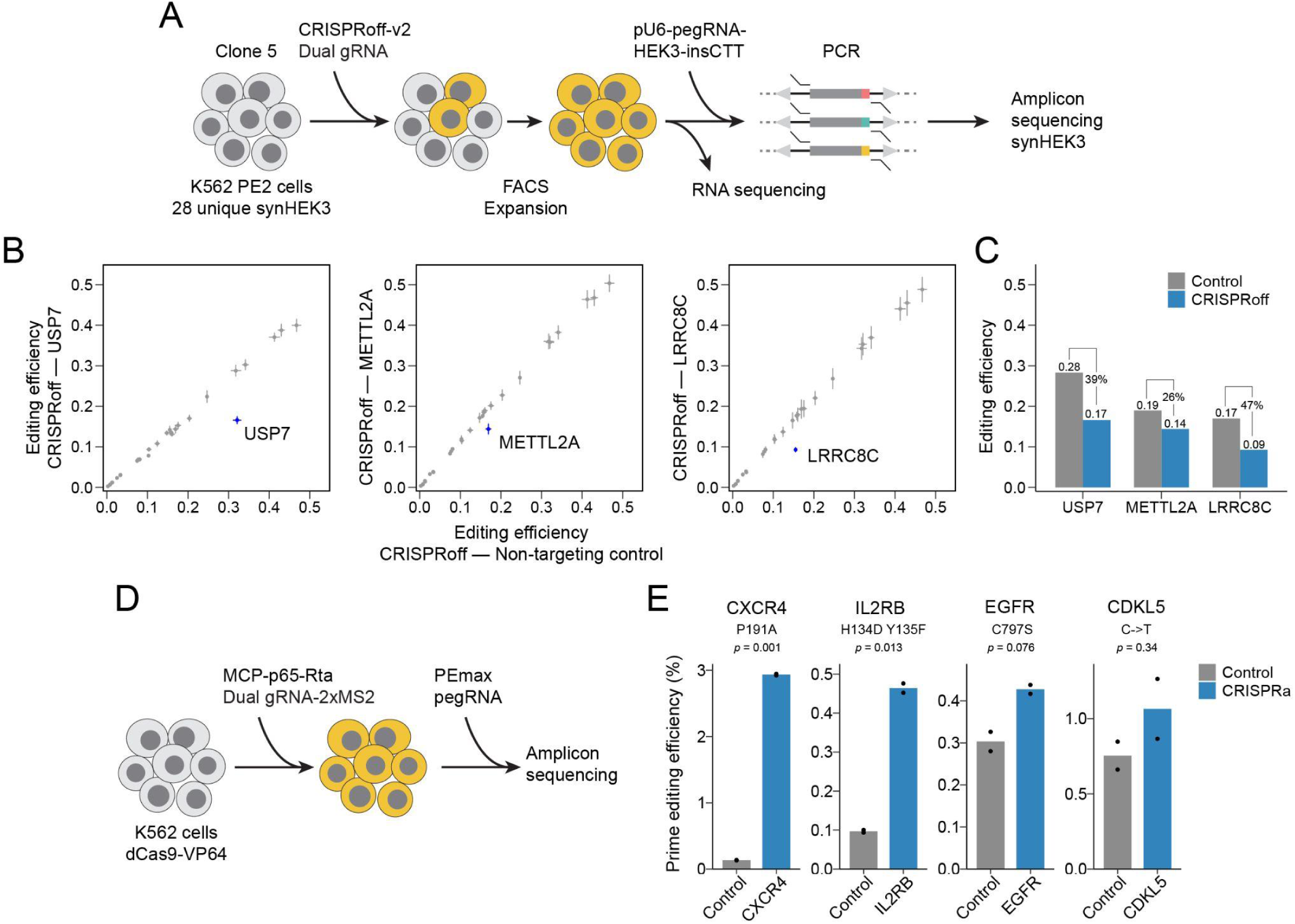
Modulating prime editing outcomes by targeted epigenetic reprogramming. **A**) Workflow of the CRISPRoff experiment. Clone 5 was transfected with CRISPRoff-v2 and paired gRNAs targeting promoters. Transfected cells were sorted on Day 2 and expanded. On Day 11, cells were transfected with a pegRNA to induce CTT insertions in synHEK3 reporters. Editing efficiencies were measured by amplicon-sequencing. The same cells were subjected to bulk RNA sequencing. **B**) Scatter plots of mean prime editing efficiencies of synHEK3 reporters in cells transfected with CRISPRoff gRNAs targeting the *USP7*, *METTL2A* and *LRRC8C* promoters (two biological replicates per condition). SynHEK3 reporters in the corresponding target genes are highlighted. Error bars correspond to standard deviation of measured editing efficiencies. **C**) Bar plot of prime editing efficiency changes of the target synHEK3 reporters in the CRISPRoff experiment. Control editing efficiencies are predicted efficiencies using linear models trained in all other synHEK3 reporters that are not in the corresponding CRISPRoff target genes as shown in panel B. Percent decreases of editing efficiencies are labeled. **D**) Workflow of the CRISPRa experiment. On Day 0, the K562 dCas9-VP64 line was transfected with paired gRNAs with two MS2 hairpins targeting gene promoters and a plasmid expressing the MCP-p65-Rta fusion. On Day 2, the cells were transfected with plasmids expressing PEmax and pegRNAs encoding specific mutations at each endogenous locus. Editing efficiencies were measured by amplicon sequencing on Day 5. **E**) Prime editing efficiencies in percentage at endogenous gene targets. *P* value: welch’s two sample t-test.

Encouraged by these results, we next examined the possibility of increasing prime editing efficiency via transient gene activation. Furthermore, for these experiments, we sought to boost the efficiency of introducing mutations to the native genome, rather than to integrated sensors. We selected four endogenous gene targets, *CXCR4*, *IL2RB*, *EGFR* and *CDKL5*, and used prime editing to model a different substitution event per target, including mutations of clinical and research importance^53–55^. For gene activation, we chose the CRISPRa Synergistic Activation Mediator (SAM) system^56^. On Day 0, we transfected a K562 cell line stably expressing the dCas9-VP64 fusion^57^ with a pair of gRNAs (2XMS2) targeting the promoter of one of these four genes, along with an MCP-p65-Rta fusion protein. On Day 2, we transfected the cells with PEmax and pegRNAs to introduce the desired mutations (**Fig. 6D**). As a result, we observed 40% (*EGFR*, *p* = 0.076) to 21.5-fold (*CXCR4*, *p* = 0.001) increases in editing frequencies when the corresponding target genes were activated by CRISPRa (**Fig. 6E**). The variation in editing frequency enhancement might be impacted by variation in the performance of CRISPRa and/or the absolute transcriptional activity at the loci targeted **Supplementary Fig. 7C,D**). However, more examples will be necessary to draw general conclusions.

Taken together, these experiments reinforce the strong link between transcriptional activity and prime editing outcome, and more importantly, demonstrate the feasibility of enhancing prime editing outcomes via epigenetic reprogramming of the gene to which edits are being introduced.

## DISCUSSION

In this study, we developed and applied a series of techniques to reveal and dissect a complex, heterogeneous influence of chromatin landscape on the efficiency of prime editing. We identified key epigenetic factors that correlated with high editing efficiency, and integrated these features to generate a predictive model for highly effective prime editing. Furthermore, by applying a pooled shRNA screen in single cells, we discovered DNA repair factors that worked in a chromatin context-specific manner. Finally, exploiting the potential causality underlying the relationship between transcriptional activity and prime editing efficiency, we showed that the efficiency of prime editing can be modulated via epigenetic reprogramming.

First, we developed a T7-assisted reporter mapping method, which proved an effective and robust framework for measuring position-dependent regulatory effects with densely integrated reporters. Using this method, we were able to saturate the integration site mapping of ∼10,000 reporters in a finite cell population (∼500 clones and 0.5-1 ug gDNA) with 1 x 10^7^ raw sequencing reads (**Supplementary Fig. 1E**). In addition to increased scale, the comprehensiveness of this method is critical for mapping *piggyBac* transposons. Due to the insertion and excision activity of the *piggyBac* transposase, 17% of the synHEK3 barcodes were found to be associated with more than one genomic location, which corresponded to 55% of all identified insertion sites. Incomplete profiling might result in incomplete removal of duplicates and noise. Of note, implementing this method only requires the addition of a 19-bp T7 promoter to any construct; therefore, it can be adapted for mapping constructs integrated via other methods, such as lentivirus, retrovirus, as well as recently characterized serine integrases^58^.

Second, the incorporation of T7 IST into contemporary single cell RNA-seq workflow enables co-profiling of non-transcribing constructs in the genome, which is critical for fully understanding phenotypes that are modulated by the local epigenetic environment, such as DNA damage response. When combined with a pooled genetic screen, this framework can be generalized for characterizing the crosstalk between cellular states, local chromatin state and any molecular processes that can be read out via a primary sequence change or barcode expression levels. We successfully applied this approach in identifying site-specific regulators of prime editing-mediated insertions. During the preparation of this manuscript, a parallel study led by Vergara *et al.* was published, that sought to determine chromatin context-dependent regulators of DNA DSB repair, using a well-based screening approach^59^. Compared to this approach, the single-cell screen strategy we developed is more flexible, can be easily adapted for use with new perturbation libraries, and is theoretically more scalable as feature-capture efficiency further increases^60^. Although the genetic screen repurposes the current single cell sequencing framework, it does not strictly require the cell’s transcriptome to be captured or sequenced. Therefore, combinatorial indexing could in principle enable the impact of vastly more perturbations (and combinations of perturbations, etc.) to be profiled, provided that the readout can be rendered compact, as it was here with the barcoded, editable synHEK3 sensor. Finally, boosting RNA abundance by post-fixation IST unlocks many new possibilities, including retrieving lineage tracing constructs in complex cellular systems and enhancing capture efficiency of feature barcodes, especially in the setting of sci-RNA-seq3.

Third, the pooled shRNA screen in 50 synHEK3 reporters uncovered a previously undescribed role of HLTF in modulating prime editing-mediated insertions, and perhaps more importantly, demonstrated the interaction between *trans*-acting DDR factors and the *cis*-chromatin environment in regulating SSB repair. Specifically, we found that HLTF preferentially suppressed prime editing in actively transcribed regions. While this phenomenon could be attributed to an unknown role of HLTF, considering its characterized functions in post-replication repair with PCNA^52, 61^, we hypothesize that HLTF might be unevenly distributed along chromatin and that actively transcribed regions of the genome might be under intense scrutinization by this protein such that DNA lesions can be corrected immediately by DNA repair machinery to avoid accumulation of deleterious mutations in the transcribed genome over cell cycles.

Fourth, current state-of-the-art technologies for improving prime editing efficiency mainly rely on enhancing the robustness of the prime editing core complex^10, 11, 13–15^, machine learning-based prediction of target site efficiencies^10, 14, 15^, and transient suppression of the MMR pathway^11, 12^. While these approaches have yielded encouraging results (especially when used in combination), there are notable limitations. Practically, the use of a codon- and structure-optimized prime editing protein or a structurally enhanced pegRNA alone produces limited effects (3-4 fold)^11, 13^. In the scenario of installing small protein tags, sequences with high insertion efficiencies predicted by a current machine learning model are only 1.63-fold better than those with low predicted efficiencies^15^. Finally, the strength of the MMR inhibition effects varies depending on the type of edit, with greater effects observed in cases of single-nucleotide substitutions and small indels^11^. Here, the new approach we developed – coupling epigenetic reprogramming with prime editing, exploits orthogonal molecular mechanisms that govern prime editing outcomes and demonstrates a new route of enhancing prime editing efficiency in addition to the approaches above. Our data show that the epigenetic environment surrounding a target can strongly impact prime editing outcome over many orders of magnitude, and transient gene activation using CRISPRa can lead to up to 21-fold increase in editing efficiency. We envision this strategy will be especially beneficial for stem cell-based gene therapy, in which the target gene remains transcriptionally silenced and is only expressed in differentiated cell types. While a bottleneck for this approach might be the efficacy of currently available gene activation strategies, as the technologies for epigenetic reprogramming continue to rapidly evolve, we anticipate that epigenetic reprogramming in order to enhance prime editing outcomes will be a highly useful and straightforward-to-implement approach.

Despite the advances in experimental strategies and our understanding of the epigenetic regulation of prime editing, a number of limitations remain to be addressed. First of all, epigenetic annotations of the synHEK3 reporters are still incomplete, which is due to the lack of epigenetic markers to better resolve heterochromatin features in K562 cells, and the lack of metrics that measure the impact of higher-order structures (*e.g.* chromatin looping) on chromatin accessibility and gene expression. This is exemplified by the highly variable editing efficiencies for synHEK3 reporters falling in distal intergenic regions (**Supplementary Fig. 2D**). Second, while the synHEK3 reporter assay has revealed a negative correlation between distance-to-TSS and prime editing efficiency, current sampling density is still far below the resolution that enables assessing the impact of individual nucleosomes on prime editing outcomes in the regions immediately surrounding TSSs, where nucleosomes are well-phased and have been shown to interfere with Cas9 binding^21, 62^. Independent strategies need to be developed to densely sample promoter-proximal regions. Third, our study only characterized prime editing to introduce 3- and 6-bp insertions at the HEK3 target site, neglecting potential interactions between target sequence, mutation type and the *cis*-chromatin environment. Future studies aimed at resolving the target-mutation-location specificities are needed to fully exploit the potential of this general approach. Fourth, in the shRNA screen with sci-RNA-seq3, the recovery rates of synHEK3 barcodes and shRNAs remain suboptimal (**Supplementary Fig. 5B**). For synHEK3, barcode recovery rate ranged from 5% to 60% and roughly correlated with their native chromatin accessibility, indicating the IST condition needs to be further optimized to overcome the physical hindrance presented by fixed chromatin. Although recent advances have shown that this task is tractable^60^, more work is in demand to achieve higher recovery of feature barcodes in sci-RNA-seq3 experiments, which critically underlies the scalability of this method. Finally, in the CRISPRa experiments, to achieve enduring activation of target genes, we utilized a cell line with integrated dCas9-VP64^57^, which could potentially compete with the prime editor for pegRNA binding, resulting in reduced concentrations of functional pegRNA and physical hindrance at prime editing target sites. Alternative strategies such as using *in vitro* assembled prime editing RNPs or orthogonal CRISPRa systems (*e.g.* saCas9-VP64) might increase the orthogonality between the systems and ameliorate the competition^63, 64^.

In conclusion, the methods and data generated in this study reveal the tight regulation of prime editing outcomes by the *cis-*chromatin landscape. Our studies also show how epigenetic reprogramming can be leveraged to enhance the efficiency of prime editing in a target-specific manner. As this strategy is orthogonal to other avenues for improving prime editing efficiency, it may be useful in both basic research and therapeutic contexts.

## SUPPLEMENTAL INFORMATION

Table S1. List of oligos and nucleic acid sequences used in this study.

Table S2. List of ENCODE datasets used in this study.

Table S3. List of synHEK3 reporters and relevant features.

Table S4. List of synHEK3 reporters mapped in the two monoclonal lines.

Table S5. The DDR-focused shRNA library.

Table S6. Processed screen results (clone 3).

Table S7. Processed screen results (clone 5).

Table S8. Prime editing efficiency measured in the CRISPRoff and CRISPRa experiments.

## Acknowledgments

We thank members of the Shendure Lab, as well as Yi Yin (UCLA), for ongoing input and advice over the course of this project. This research is supported by research grants from the National Human Genome Research Institute (NHGRI; UM1HG011966 to J.S., R01HG010632 to J.S.). D.C. was supported by award no. F32HG011817 from the National Human Genome Research Institute. J.C. is a Howard Hughes Medical Institute Fellow of the Damon Runyon Cancer Research Foundation (DRG-2403-20). T.A.M was supported by a Banting Postdoctoral Fellowship from the Natural Sciences and Engineering Research Council of Canada (NSERC). J.B.L. is a Fellow of the Damon Runyon Cancer Research Foundation (DRG-2435-21). J.F.N is supported by the UW MSTP Precision Medicine Fellowship. J.S. is an Investigator of the Howard Hughes Medical Institute.

## Author contributions

Conceptualization, X.L., W.C. and J.S.; Methodology, X.L., W.C., B.K.M. and J.S.; Investigation: X.L.; Formal Analysis and Software, X.L.; Writing, X.L. and J.S.; Funding Acquisition, J.S.; Resources: D.C., C.L., J.C., F.M.C., T.A.M., H.K., J.B.L. and J.F.N; Supervision, J.S.

## RESOURCE AVAILABILITY

### Materials Availability

All unique reagents generated in this study are available from the Lead Contact with a completed Materials Transfer Agreement.

### Data and Code Availability

The accession number for the datasets reported in this paper is GEO: GSE228465. Code and scripts are being organized and will be released soon.

## EXPERIMENTAL MODEL AND SUBJECT DETAILS

### Cell culture conditions

K562 cells (CCL-243) were purchased from ATCC and maintained in RPMI 1640 medium (Gibco) supplied with 10% FBS (Hyclone) and penicillin/streptomycin (Gibco, 100 U/ mL). HEK293T cells were maintained in DMEM medium (Gibco) supplied with 10% FBS and penicillin/streptomycin. All cells were kept in a humidified incubator at 37 °C, 5% CO_2_.

## METHOD DETAILS

### Molecular cloning

#### PB-T7-HEK3-BC (synHEK3)

First, a minimal *piggyBac* cargo construct was created by deleting all intervening sequences between the 5’ and 3’ terminal repeats (including core insulators) of the PB-CMV-MCS-EF1α-Puro vector (System Biosciences, PB510B-1). A gBlock (Integrated DNA Technologies, IDT) consisting of a filler sequence and flanking scaffold sequences (from GFP) was inserted to create a shuttle vector. The filler sequence contains two divergent BsmBI recognition sites and can be removed scarlessly. Second, the shuttle vector was digested with BsmBI (New England Biolabs). A 87-bp region around the HEK3 gRNA target site was synthesized from IDT and amplified with a pair of primers to introduce a T7 promoter and a 16-bp barcode to its upstream and downstream, respectively. The resulting PCR product was inserted into the linearized shuttle vector using NEB HiFi assembly. Ligated products were cleaned up and concentrated with AMPure XP beads (Beckman Coulter), and electroporated into NEB 10-beta electrocompetent *E. coli*. Electroporation was performed in a 0.1 cm electroporation cuvette using a Bio-Rad GenePulser electroporator at 2.0 kV, 200 Omega, and 25 μF.

#### LT3-GFP-T7-miR-E-CS1-PGK-Neomycin

The LT3GEN vector was purchased from Addgene (#111173) and digested with I-SceI (New England Biolabs). A fragment containing a T7 promoter and homologous sequences was ordered from IDT and assembled into the backbone. The LT3-GFP-T7-PGK-Neomycin vector was digested with XhoI and EcoRI-HF (New England Biolabs). An shRNA targeting the Renilla luciferase (Ren713) was inserted along with a Capture Sequence 1 (CS1) after the EcoRI site. For shRNA cloning, the LT3-GFP-T7-miR-E-CS1-PGK-Neomycin construct was digested with XhoI and EcoRI-HF. shRNAs were ordered from IDT as 97-nt 4 nmole Ultramers or oPool and amplified with primers p1 (5’-ATTACTTCGACTTCTTAACCCAACAGAAGGCTCGAGAAGGTATATTGCTGTTGACAGTGAGCG-3’) and p2 (5’-AATTGCTCTTGCTAGGACCGGCCTTAAAGCGAATTCTAGCCCCTTGAAGTCCGAGGCAGTAGGCA-3’). The PCR products were assembled into the backbone using NEB HiFi assembly and transformed into NEB Stable Competent *E. coli* (single shRNA vectors) or electroporated into NEB 10-beta electrocompetent *E. coli* (library).

#### Lenti-rtTA-P2A-Blast

The Lenti-Cas9-P2A-Blast (Addgene: 52962) vector was digested with XbaI and BamHI (New England Biolabs) to create a backbone. rtTA was amplified from the LT3GEPIR vector (Addgene: 111177) and cloned into the backbone using NEB HiFi assembly.

#### pU6-Sp-pegRNA-HEK3-ins6N and pU6-Sp-pegRNA-HEK3-ins6N

the pU6-Sp-pegRNA-HEK3-insCTT vector (Addgene:132778)^1^ was linearized by PCR with 5’ phosphorylated oligos p3 (5’-TCTGCCATCANNNNNNCGTGCTCAGTCTGTTTTTTTAAGCTTG-3’, ins6N) or p4 (5’-TCTGCCATCANNNCGTGCTCAGTCTGTTTTTTTAAGCTTG-3’, ins3N) and p5 (5’-GGACCGACTCGGTCCCACTT-3’) and ligated with T4 DNA ligase. Ligation product was concentrated and electroporated into NEB 10-beta electrocompetent *E. coli*.

#### pU6-Sp-dual-gRNA vectors

A pU6-Sp-dual-gRNA scaffold vector was generated by replacing the pegRNA expressing cassette of pU6-Sp-pegRNA-HEK3-insCTT vector with a dual U6-gRNA cassette from the PX333 vector (Addgene: 64073)^65^. The second gRNA cloning site (two BsaI sites) was modified to two BsmBI sites. Spacer sequences were cloned into this vector sequentially using the oligo annealing method^66^.

#### pU6-Sp-gRNA-2XMS2 vectors

A pU6-Sp-gRNA-2XMS2 scaffold vector was generated by modifying the pU6-Sp-pegRNA-HEK3-insCTT backbone. The scaffold sequence is 5’-GTTTAAGAGCTAAGCCAACATGAGGATCACCCATGTCTGCAGGGCATAGCAAGTTTAAATAAGGCTAGTCCGTTATCAACTTGGCCAACATGAGGATCACCCATGTCTGCAGGGCCAAGTGGCACCGAGTCGGTGCTTTTTTT-3’^28^. Spacer sequences were cloned in between two BsmBI sites using the oligo annealing method^66^.

#### PB-CMV-MCP-XTEN80-p65-Rta-3xNLS-P2A-T2A-mPlum

The MCP(N55K) sequence^67^ was synthesized as an IDT gBlock and amplified. XTEN80 and 3XNLS-P2A were amplified from TETv4 (Addgene: 167983)^28^. p65-Rta was amplified from sadCas9-VPR (Addgene: 188514)^68^. mPlum was amplified from mPlum-C1(Addgene: 54839)^69^. All PCR products were purified and cloned into the PB-CMV-MCS-EF1α-Puro vector between the NotI site and the SV40 polyA sequence using NEB HiFi assembly. Then, a T2A sequence was inserted next to P2A by NEB HiFi assembly.

pU6-Sp-pegRNA plasmids used in this study were ordered as 4nm ultramers (IDT) or long primers containing the spacer and 3’ extension sequences and cloned into the backbone of pU6-Sp-pegRNA-HEK3-CTT using NEB HiFi assembly.

### Transfection strategies

Transfections of K562 cells were performed using a Lonza Bioscience 4D-Nucleofector system and the SF Cell Line 4D-Nucleofector X kits (Lonza). For single nucleocuvettes (100 uL), 1-2 x 10^6^ cells were transfected with up to 4 ug DNA. For 96-well Nucleocuvette plates (20 uL), 2 x 10^5^ ∼ 5 x 10^5^ cells were transfected with up to 2 ug DNA. Program code was FF-120.

#### CRISPRof f experiments

On Day 0, 2 x 10^6^ clone 5 cells were transfected with CRISPRoff-v2.1 (3,000 ng), pU6-Sp-dual-gRNA (1,000 ng) and pmax-GFP (500 ng, Lonza) using the SF Cell Line 4D-Nucleofector X kit L (Lonza). On Day 2, cells were sorted based on high GFP expression (top 20%) on a flow cytometer and expanded. On Day 11, 4 x 10^5^ cells were transfected with the pU6-Sp-pegRNA-HEK3-insCTT plasmid (1,000 ng) using the SF Cell Line 96-well Nucleofector Kit (Lonza). Program code was FF-120.

#### CRISPRa experiments

On Day 0, 4 x 10^5^ K562 dCas9-VP64 cells were transfected with PB-CMV-MCP-XTEN80-p65-Rta-3xNLS-P2A-T2A-mPlum (600 ng) and paired pU6-Sp-gRNA-2XMS2 (200 ng each) plasmids targeting the same promoter. On Day 2, 4 x 10^5^ cells from the previous transfection were transfected with pCMV-PEmax (Addgene: 174820)^11^ (800 ng) and the pU6-Sp-pegRNA plasmids (400 ng). All transfections were performed using the SF Cell Line 96-well Nucleofector Kit (Lonza) with program FF-120.

For Cas9 RNP transfection, Alt-R sgRNAs were ordered from IDT and resuspended in TE buffer to 100 uM. The RNP complex was assembled by combining 2.1 uL PBS, 1.2 uL Alt-R sgRNA (100 uM) and 1.7 uL Alt-R Cas9 Nuclease (61 uM, IDT) and incubating at room temperature for 10-20 min. The RNP mixture was added to cells resuspended in nucleofection solution along with 0.5 ug pmax-GFP (Lonza) and 1 uL electroporation enhancer (100 uM, IDT). Nucleofection was performed using the program FF-120.

Transfections of the HEK293T cell line were performed using Lipofectamine 3000 (Invitrogen) following manufacturer’s instructions.

### Lentivirus production and transduction

HEK293T cells (1.6 x 10^7^) were seeded in a 10 cm^2^ dish the day before transfection. 5.6 ug lentiviral vector and 14.4 ug ViraPower Lentiviral Packaging Mix (Invitrogen) were mixed and transfected using Lipofectamine 3000 (Invitrogen) following manufacturer’s instructions. Medium was changed at 24 hours post transfection. Viruses were collected at 48 and 72 hours post transfection, filtered using 45 um filters and concentrated 100-fold using PEG-it Virus Precipitation Solution (System Biosciences).

In general, K562 cells were transduced with lentivirus in the presence of 8 ug/mL polybrene (Millipore). Medium was replaced after 24 hours. Clones 3 and 5 were transduced with Lenti-rtTA-P2A-Blast viruses and selected in 10 ug/mL Blasticidin (Invitrogen) for 7 days. Monoclonal lines were derived for the downstream shRNA screen. To generate high-MOI transduced K562 cells for the shRNA screen, the lentiviral library was first titrated. Then, cells were mixed with an adjusted amount of virus aiming for an MOI of 10. Transduced cells were selected in 800 ug/mL Geneticin (Invitrogen) for 7 days.

### Quantitative PCR (qPCR) analysis

qPCRs were performed on purified gDNA or cDNA reverse transcribed from total RNAs using SuperScript IV reverse transcriptase (200 U/uL, Invitrogen) following manufacturer’s instructions. qPCRs were performed with Power SYBR Green PCR Master Mix (Invitrogen) or KAPA2G Robust HotStart ReadyMix (Roche) supplied with SYBR Green (Invitrogen). For copy number estimation of synHEK3 reporters, Cq values of synHEK3 were normalized to those of SNRPB (3 copies per genome). See **Table S1** for the list of primers used.

### T7-assisted reporter mapping

gDNA of K562 cells was purified using the DNeasy Blood & Tissue Kit (QIAGEN) and *in vitro* transcribed with the HiScribe T7 High Yield RNA Synthesis Kit (New England Biolabs). Briefly, each reaction (20 uL) contained 0.3∼1 ug gDNA, NTPs (10 mM each) and 2 ul T7 RNA Polymerase Mix. The reaction mixture was incubated at 37 °C for 16 hours. Then, gDNA was digested with 2.5 uL DNase (QIAGEN) in a 100 uL reaction at room temperature for 30 min. RNA was extracted with TRIzol LS Reagent (Invitrogen), and aqueous phase was precipitated with 1 volume of isopropanol and 5 ug Glycogen (Invitrogen) at -80 °C for 1 hour. RNA pellet was collected by centrifugation at 21,000 x g at 4 °C for 1 hour. The pellet was washed with 80% ice-cold ethanol and resuspended in 11.5 uL nuclease-free water.

For reverse transcription, RNA was incubated with 0.5 uL 100 uM RT primer p6 (5’-TCGTCGGCAGCGTCAGATGTGTATAAGAGACAGNNNNNNNN-3’) and 1 uL 10 mM dNTP at 65 °C for 5 min and cooled on ice. Then, 4 uL 5X RT buffer, 1 uL 100 mM DTT, 1 uL RNaseOUT (40 U/uL) and 1 uL SuperScript IV RT (200 U/uL, Invitrogen) were added and the reaction mixture was incubated at 23 °C for 10 min, 50 °C for 15 min and 80 °C for 10 min.

cDNA library was amplified with KAPA2G Robust HotStart polymerase, using primers p7 (5’- GTGACTGGAGTTCAGACGTGTGCTCTTCCGATCTGAAAGGAAGCCCTGCTTCCTCCAGAGGG-3’, 0.5 uM) and p8 (5’- TCGTCGGCAGCGTCAGATGTGTATAAGAGACAG-3’, 0.5 uM). PCR reaction was performed as follows: 95 °C 3 min; 95 °C 15 s, 65 °C 15 s, 72 °C 30 s, 16∼18 cycles; and 72 °C 1 min. The PCR product was subjected to double-sided size selection (0.5X, 0.9X) and cleaned up with AMPure XP beads. The resulting product ranged from 200 to 1000 bp. To prepare Illumina sequencing libraries, 5-10 ng PCR product was re-amplified with the Nextera P5 and TruSeq P7 library index primers as shown in **Table S1** for 5 cycles. The final PCR product underwent another round of double-sided size selection (0.5X, 0.9X) and cleaned up with AMPure XP beads. The library was sequenced on an Illumina MiSeq in paired-end mode (Read 1: 254 bp; Read2: 55bp).

### HEK3 editing library preparation

gDNA of K562 cells was purified using the DNeasy Blood & Tissue Kit. Alternatively, cells were lysed in a lysis buffer [100 mM Tris-HCl pH8.0, 0.05% SDS and 0.04 mg/mL proteinase K(Thermo Scientific)], and incubated at 50 °C 60 min and 80 °C 30 min. 100∼250 ng gDNA or cell lysates were amplified using KAPA2G Robust HotStart ReadyMix with primers p9 (5’-GTGACTGGAGTTCAGACGTGTGCTCTTCCGATCTCTACCCCGACCACATGAAGCAGC-3’, 0.5 uM) and p10 (5’-TCGTCGGCAGCGTCAGATGTGTATAAGAGACAGNNNNNNNNNNGACCATGTCATCGCGCTTCTCGT-3’, 0.5 uM). PCR reaction was performed as follows: 95 °C 3 min; 95 °C 15 s, 68-N °C 15 s, 72 °C 30 s, for 9 cycles (N was cycle number); 95 °C 15 s, 65 °C 15 s, 72 °C 30 s, for 11 cycles; and 72 °C 1 min. To ensure enough coverage and accurate measurement of editing efficiencies, for the K562 synHEK3 pool, we pooled products from at least 16 PCR reactions. The PCR product was purified with AMPure XP beads. 5 ng PCR product was re-amplified with the Nextera P5 and TruSeq P7 library index primers in **Table S1** for 5 cycles. The final libraries were cleaned up with AMPure XP beads and sequenced on an Illumina NextSeq 500 or an Illumina NextSeq 2000 sequencer.

### *In situ* transcription and sci-RNA-seq3 library preparation

sci-RNA-seq3 libraries were prepared by modifying a recently published protocol from the lab^26^. Briefly, K562 cells were collected, counted, washed with PBS and lysed in 5 mL Hypotonic lysis buffer B with DPEC (Sigma) for 10 min on ice. Nuclei were collected by centrifugation at 500 xg, 4 °C for 3 min and resuspended in 1 mL 0.3 M SPBSTM buffer with DEPC. Nuclei were fixed with 4 mL ice-cold methanol for 15 min on ice, swirled occasionally. After rehydration by adding 10 mL SPBSTM, nuclei were collected by centrifugation and washed once with 1 mL 0.3 M SPBSTM. For T7 *in situ* transcription, fixed nuclei resuspended in 171 uL SPBSTM were mixed with 99 uL NTP buffer and 30 uL T7 polymerase mix (New England Biolabs) and incubated in a 1.5 mL LoBind tube (Eppendorf) at 37 °C for 30 min. Afterwards, nuclei were washed with 1 mL SPBSTM, sonicated using a Diagenode sonicator for 12 seconds. At this point, nuclei were stained with YOYO-1 Iodide (Invitrogen) and counted on a Countess automated cell counter.

Counted nuclei were resuspended in SPBSTM at 4 x 10^6^/mL. For each RT plate, 500 uL nuclei were combined with 56 uL 10 mM dNTPs (New England Biolabs) and distributed to a low-bind 96-well plate (5 uL per well). 1 uL indexed oligo-dT, HEK3 and CS1 primers (10 uM) were added to each well. The plate was incubated at 55 °C for 5 min and cooled on ice. RT mixture was prepared by combining 240 uL 5X SuperScript IV Buffer, 60 uL SuperScript IV and 60 uL water. 3 uL RT mixture was added to each well. The RT plate was incubated at 55 °C for 10 min and cooled on ice. 5 uL ice-cold SPBSTM was added to each well and all nuclei were pooled in pre-chilled LoBind tubes. Nuclei were washed once with 1 mL SPBSTM and resuspended in 1,200 uL SPBSTM (per ligation plate).

11 uL nuclei were distributed to each well of a new 96-well plate and mixed with 2 uL 10 uM ligation primers. For each ligation plate, 195 uL 10X T4 Ligation Buffer was mixed with 65 uL T4 DNA Ligase, and 2 uL of the ligation mixture was added to each well. Ligation was performed at room temperature for 20 min on the bench. For the shRNA screen, nuclei were distributed into 4 ligation plates (384 ligation indices) to increase cell index complexity. The ligation plate was then cooled on ice and 10 uL ice-cold SPBSTM was added to pool nuclei from all wells. Nuclei were washed twice with 1 mL SPBSTM.

Nuclei were resuspended in 1X Second Strand Synthesis Buffer (New England Biolabs), counted with YOYO-1 Iodide and diluted to 1.3 x 10^5^ - 2.5 x 10^5^ per 400 uL. 4 uL nuclei were distributed to multiple 96-well plates based on the number of nuclei retrieved at this step. Extra plates with nuclei were stored at -80 °C. Second strand synthesis mixture was prepared by combining 10.5 uL 10X Second Strand Synthesis Buffer, 35 uL 20X Second Strand Synthesis Enzyme with 94.5 uL water. 1 uL of second strand synthesis mixture was added to each well. The plate was incubated at 16 °C for 2.5 hours. To lyse nuclei, 1 uL ∼1.07 AU/mL Protease (Qiagen) was added to each well and incubated at 37 °C for 40 min. Protease was inactivated at 75 °C for 20 min. 5 uL EB buffer (QIAGEN) was added to each well and mixed. 5 uL was taken out from each well and used for enrichment PCR, while the rest was used for Tn5 tagmentation and transcriptome library preparation.

To prepare the transcriptomic library, Tn5-N7 and Mosaic end oligos were resuspended to 100 uM in annealing buffer (50 mM NaCl, 40 mM Tris-HCl pH8.0), mixed at a 1:1 ratio and annealed on a thermocycler using the following program: 95°C 5 min, cool to 65°C (0.1°C/s), 65°C 5 min, cool to 4°C (0.1°C/s). 20 uL Tagmentase (Tn5 transposase - unloaded, Diagenode) was mixed with 20 uL annealed oligos and incubated on a thermomixer at 350 rpm, 23°C for 30 min. 20 uL glycerol was added to the loaded Tn5 before storage at -20 °C. For tagmentation, 13.75 uL N7-loaded Tn5 (Diagenode) was mixed with 550 uL TD buffer and 5 uL of this mixture was added to each well. The plate was incubated at 55 °C for 5 min. To remove Tn5 transposase, 50 uL 1% SDS, 50 uL BSA (New England Biolabs) and 225 uL water were mixed, and 2.6 uL of the mixture was added to each well and incubated at 55 °C for 15 min. Then, SDS was quenched by adding 2 uL 10% Tween-20 to each well. For PCR, 96 indexed P5 primers were used with constant or indexed P7 primers (See **Table S1**). A PCR master mixture was prepared by combining 2,200 uL 2x NEBNext High-Fidelity 2X PCR Master Mix (New England Biolabs), 22 uL common P7 primer (100 uM) and 352 uL water. 2 uL indexed TruSeq P5 primer and 23.4 uL PCR mixture were added to each well. If using indexed P7 primers, 2 uL of 10 uM primer was added to each well. PCR reaction was performed as follows: 70 °C 3 min; 98 °C 30 s; 98 °C 10 s, 63 °C 30 s, 72 °C 60 s, 16 cycles; and 72 °C 5 min. 3 uL of each well was pooled and cleaned up with 0.8X AMPure XP beads. The library was resolved on a 1% agarose gel and the smear between 300-600 bp was extracted using Monarch DNA Gel Extraction Kit (New England Biolabs).

For HEK3 and shRNA libraries, enrichment PCRs were directly performed on Protease-treated cDNAs. To match PCR indices with those of the transcriptome library, the same indexed TruSeq P5 primers were used. The enrichment PCR master mixture contained 2,200 OneTaq 2X Master Mix (New England Biolabs), 16.5 uL synHEK3 P7 enrichment primer (100 uM), 16.5 uL shRNA P7 enrichment primer (100 uM), 44 uL 100X SYBR green (Invitrogen) and 1,353 uL water. 2 uL indexed P5 primer and 33 uL PCR reaction was performed as follows: 95 °C 3 min; 95 °C 15 s, 68-N °C 15 s, 72 °C 30 s, for 9 cycles (N was cycle number); 95 °C 15 s, 65 °C 15 s, 72 °C 30 s, for M cycles (decided by qPCR); and 72 °C 1 min. Monitor the reaction on a real-time qPCR machine and terminate the reaction at the log phase. All PCR products were pooled and concentrated using 0.9X AMPure XP beads. The library was resolved on a 1% agarose gel and the two discrete bands corresponding to HEK3 and shRNA constructs were extracted using Monarch DNA Gel Extraction Kit.

Eventually, all libraries were sequenced on an Illumina NextSeq 500 or an Illumina NextSeq 2000 sequencer. For HEK3 and shRNA libraries, custom Index 1 primer and Read 2 primer were used. Usually, 34 cycles were allocated to Read 1 for reading the RT index, ligation index, and unique molecular identifier (UMI). See **Table S1** for a list of oligos and sequencing primers used in this experiment.

### Amplicon sequencing for endogenous genes

Cells were lysed in a lysis buffer (100 mM Tris-HCl pH8.0, 0.05% SDS and 0.04 mg/mL proteinase K), and incubated at 50 °C 60 min and 80 °C 30 min. Lysates were directly used for PCR with the KAPA2G Robust HotStart ReadyMix with primers (0.5 uM each) designed for the endogenous targets. PCR reaction was performed as follows: 95 °C 3 min; 95 °C 15 s, 66-N °C 15 s, 72 °C 40 s, for 8 cycles (N was cycle number); 95 °C 15 s, 60 °C 15 s, 72 °C 40 s, for 11 cycles; and 72 °C 1 min. The PCR product was purified with AMPure XP beads. 5 ng PCR product was re-amplified with the Nextera P5 and TruSeq P7 library index primers in **Table S1** for 5 cycles. The final libraries were cleaned up with AMPure XP beads and sequenced on an Illumina Miseq, an Illumina NextSeq 500 or an Illumina NextSeq 2000 sequencer.

Sequencing reads were demultiplexed using the bcl2fastq software (Illumina). A custom script was used to determine frequencies of the introduced substitutions.

### RNA sequencing library preparation and processing

For bulk RNA sequencing of K562 PE2-Puro cells, total RNAs were purified from cells using TRIzol LS Reagent (Invitrogen) following manufacturer’s instructions, treated with RNase-free DNase (QIAGEN) and cleaned up using RNeasy Mini kit (QIAGEN). 1 ug total RNA (RNA integrity >= 9.7) were used for unstranded mRNA library preparation using the TruSeq RNA Library Prep Kit v2. For the CRISPRoff experiment, total RNAs were purified from cells using TRIzol LS Reagent, treated with Turbo DNase (Invitrogen) and cleaned up using the RNeasy Mini kit. 1 ug total RNA (RNA integrity >= 9.9) were used for stranded mRNA library preparation using the Illumina TruSeq® Stranded mRNA Library Prep kit. The RNA libraries were indexed using the Illumina TruSeq RNA UD Indexes. All libraries were sequenced on an Illumina NextSeq 500 or an Illumina NextSeq 2000 sequencer in a paired end mode.

Sequencing reads were demultiplexed using the bcl2fastq software (Illumina). For calculating TPM of genes, sequencing reads from the unstranded RNA libraries were aligned to the GRCh38 reference genome (Gencode V43) using Salmon (v1.9.0)^70^. For the CRISPRoff experiment, on average, 55 million 50-bp paired-end reads were obtained per sample. Reads were aligned to the GRCh38 reference genome and counted against all Ensembl genes using STAR (2.7.6a)^71^. Raw counts were analyzed with DESeq2^72^.

### T7-assisted reporter mapping library processing

Sequencing reads were first demultiplexed using the bcl2fastq software. Under the current library design and sequencing scheme, Read1 started from genomic sequence and extended into the integrated synHEK3 construct, while Read2 contained reporter barcode information. Therefore, 1) for each sequencing read pair, the 16-bp reporter barcode was extracted from Read 2 and attached to the read name of Read 1. 2) Read1 was trimmed using cutadapt (v4.1)^73^ with following parameters: --cores=4 --discard-untrimmed -e 0.2 -m 10 -O 8 -a CCCTAGAAAGATAGTCTGCGTAAAATTGACGCATG. The adapter corresponded to the 3’ITR of *piggyBac* transposon. Since parameter “--discard-untrimmed” was used, only reads spanning insertion junctions were kept and trimmed. 3) Trimmed sequences were aligned to the GRCh38 reference genome using bwa mem (v0.7.17)^74^. The “-Y” option was used to enable soft clipping for supplementary alignments. 4) Reads uniquely (without XA:Z tag) and contiguously mapped near putative *piggyBac* landing pads (TTAA motifs) were kept using samtools (v1.9)^74^ and a custom script. 5) Aligned reads were converted to BED format using the sam2bed function in bedops (v2.4.35)^75^. Reads aligned to standard chromosomes were kept. 6) Insertion points were calculated for all reads based on strand of alignment. And reads were sorted by the insertion coordinates. 7) The first 8-bp of reads were used as UMIs. A custom script was used to collapse reads at a per-location, per-barcode, per UMI basis. A barcode-location-UMI count table was generated. 8) synHEK3 barcodes <3 Levenshtein Distance at each location were collapsed and barcodes with > 3 UMIs were kept. 9) The count table was converted to a GenomicRanges^76^ object in R. Coordinates of the last 4 base pairs of aligned reads were designated as genomic locations of the inserted synHEK3 reporters. 10) “Landing pads” of the mapped synHEK3 reporters were retrieved using the getSeq() function in the BSgenome package. Reporters that didn’t have a TTAA sequence (which could be due to PCR error, mapping error, or the use of non-canonical landing pads) were removed. 11) SynHEK3 barcodes mapped to more than one location were removed.

Aligned reads were visualized on the Integrated Genome Browser (IGV; **Fig. 1B**)^77^. Motif enrichment and visualization (**Fig. 1C**) was performed using WebLogo 3^78^. Genomic coverage analysis and annotation of the synHEK3 reporters (**Fig. 1D,E)** were performed using the ChIPseeker package^79, 80^ in R. Enrichment of synHEK3 reporters across genomic features (**Supplementary Fig. 1G,H**) were performed using the annotatePeaks.pl function in Homer^81^. A custom script was used to find overlapping genes and calculate distance to closest TSSs.

### Analysis of ENCODE datasets

Accessions of ENCODE datasets (bigwig files) used in this study are listed in **Table S2**. The GenomicRanges object containing all unique synHEK3 reporters was resized to 2 kb in width. The R package Genomation^82^ was used to extract values within the 2-kb windows from the bigwig files of the chromatin features. For a specific dataset, scores of the synHEK3 reporters were calculated as 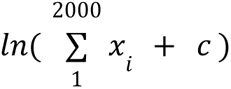, where *x*_*i*_ was the value at base *i*, and *c* was the minimum non-zero value of the epigenetic feature in a base within all the windows surveyed. For the generation of heatmap in **Fig. 5B**, chromatin feature scores were scaled using the scale() function in R. Clustering and visualization of synHEK3 reporters was performed using the ComplexHeatmap package^83^ in R.

### synHEK3 editing library analysis

Sequencing reads from the Illumina NextSeq platforms were first demultiplexed using the bcl2fastq software. The 16-bp reporter barcodes were extracted from read and attached to its read name. Sequences around the CRISPR cut site were extracted for all reads. A barcode-editing outcome table was generated. For prime editing experiments, a custom script was used to align sequences to a reference sequence and count mutation frequency for every barcode. For Cas9 mutagenesis analysis, all sequences were aggregated and editing outcomes (alleles) with the highest number of counts were selected. These most frequent alleles were then aligned to reference sequences using needleall^84, 85^ with the following parameters: -gapopen 20-gapextend 0.5 -endopen 20. A custom script was used to annotate the mutational events. MMEJ alleles were selected based on the following criteria: 1) microhomology sequences being at least 2 bp; 2) observed allele frequencies being 6-fold higher than expected frequencies. The 9 most frequent non-wild-type alleles were used for calculating the MMEJ/(MMEJ+NHEJ) ratio.

### sci-RNA-seq3 transcriptome library processing

Sequencing reads from the Illumina NextSeq platforms were first demultiplexed based on PCR index using the bcl2fastq software. Reads were filtered based on RT and ligation index, by allowing <2 Hamming distance from reference. Filtered reads were trimmed with TrimGalore (v0.6.6) with parameters: -a AAAAAAAA--three_prime_clip_R1 1. Reads were then aligned to the GRCh38 reference genome using STAR (v2.7.6a). PCR duplicates were collapsed using the RT index, ligation index, UMI sequence and end coordinate of reads. Reads were further demultiplexed based on the combination of the RT, ligation and PCR index and split into files for individual cells. To generate gene expression count matrices, reads were assigned to the exonic and intronic region of closest genes with HTseq (v.2.0.2)^86^. Reads with ambiguous assignments were discarded. Cells were further filtered based on total UMI (> 100) and mitochondrial reads percentage (<10%). Cells with the number of features detected between 10% and 90% percentile of all cells were kept and considered high-quality cells. The single cell analysis was performed using the Seurat (v4.0.0) package^87^ in R.

### Prime editing screen analysis

Sequencing reads from the Illumina NextSeq platforms were first demultiplexed based on PCR index using the bcl2fastq software. A custom script was used to further demultiplex and filter the synHEK3 and shRNA libraries based on the combination of RT, ligation and PCR index (cell ID), by allowing <2 Hamming distance from reference. For the synHEK3 library, reporter barcodes (<2 Hamming distance to the reference barcode set), prime editing outcomes and read UMIs were extracted. For the shRNA library, shRNA sequences (<2 hamming distance to the reference barcode set) and read UMIs were extracted. UMIs with <3 Hamming distance were collapsed.

A series of pre-processing steps were applied to the data. 1) We matched cell IDs between the sci-RNA-seq3 transcriptome libraries and the capture libraries, and only kept high-quality cells (number of features between 10% and 90% percentile). 2) For the shRNA library, cells with <3 or >200 UMIs or >20 shRNA were removed. 3) For the synHEK3 library, cells with <4 UMIs were first removed. We counted UMIs for synHEK3 reporters belonging to the two clones and calculated a clone UMI/total UMI ratio. If the ratio was >80% for a specific clone, the cell is assigned to the corresponding clone. Cells with extremely high UMIs (cutoff between 100 and 250 UMIs, depending on library complexity and sequencing depth of the plate) were also removed for downstream analysis. 4) For a cell-synHEK3 barcode combination, if multiple insertion sequences were observed, sequences with <3 Hamming distance were collapsed. Cell-synHEK3 barcode pairs having conflicting editing outcomes were discarded. 5) A cell-synHEK3 barcode-editing outcome-shRNA matrix was generated for a common set of cells identified by the shRNA and synHEK3 libraries.

In general, the prime editing efficiency of a specific synHEK3 reporter was estimated in all cells containing the reporter by collapsing reads on a per-cell, per-barcode basis. To assess the effect of an shRNA on a specific synHEK3 reporter, editing efficiency of this reporter was calculated in cells without the specific shRNA, which was defined as the hypothesized probability of editing. Then, the total number of cells containing the shRNA and number of cells with 6-bp insertions were counted. A *p* value was computed using the function binom.test() in R. For each clone, all synHEK3-shRNA pairs were assessed by binomial tests. The resulting *p* values were corrected using the function p.adjust() in R and the Benjamini-Hochberg method was used. Empirical FDRs of candidate synHEK3-shRNA pairs were calculated as: eFDR = (n_lower + 1)/(N + 1), where n_low was the number of control tests with a *p* value lower than the candidate test’s raw *p* value, and N was the total number of control tests. Processed screen results for clone 3 and clone 5 are provided in **Table S6** and **Table S7**.

### QUANTIFICATION AND STATISTICAL ANALYSIS

The statistical tests for each experiment are described in the text, figure legends or STAR methods. In **Fig. 2D**, the beta regression model was performed using the function betareg() in R package betareg^88^ using default parameters. In **Supplementary Fig. 2C, Supplementary Fig. 3D** and **3E**, the fit with confidence intervals was produced using function ‘geom_smoonth()’ in the R package ggplot using parameter ‘method = ‘‘lm’’’ and all other parameters being default. In **Fig. 4**, binomial tests were performed to measure the effects of shRNAs and the resulting *p* values were corrected for multiple hypothesis testing using the Benjamini & Hochberg method. In **Fig. 5C**, Fisher’s exact test was used to compare the differential enrichment of TxDb-overlapping synHEK3 reporters in the two groups. In **Fig. 5D**, *p* value was calculated using a two-sided Kolmogorov–Smirnov test. In **Supplementary Fig. 6C** and **6D**, *p* values were calculated using two-sided Kolmogorov–Smirnov tests. In **Fig. 6E** and **Supplementary Fig. 7C**, welch’s two sample t-tests were used to compare prime editing efficiency or gene expression differences of genes before and after CRISPR activation.

---

**Supplementary Figure 1.**
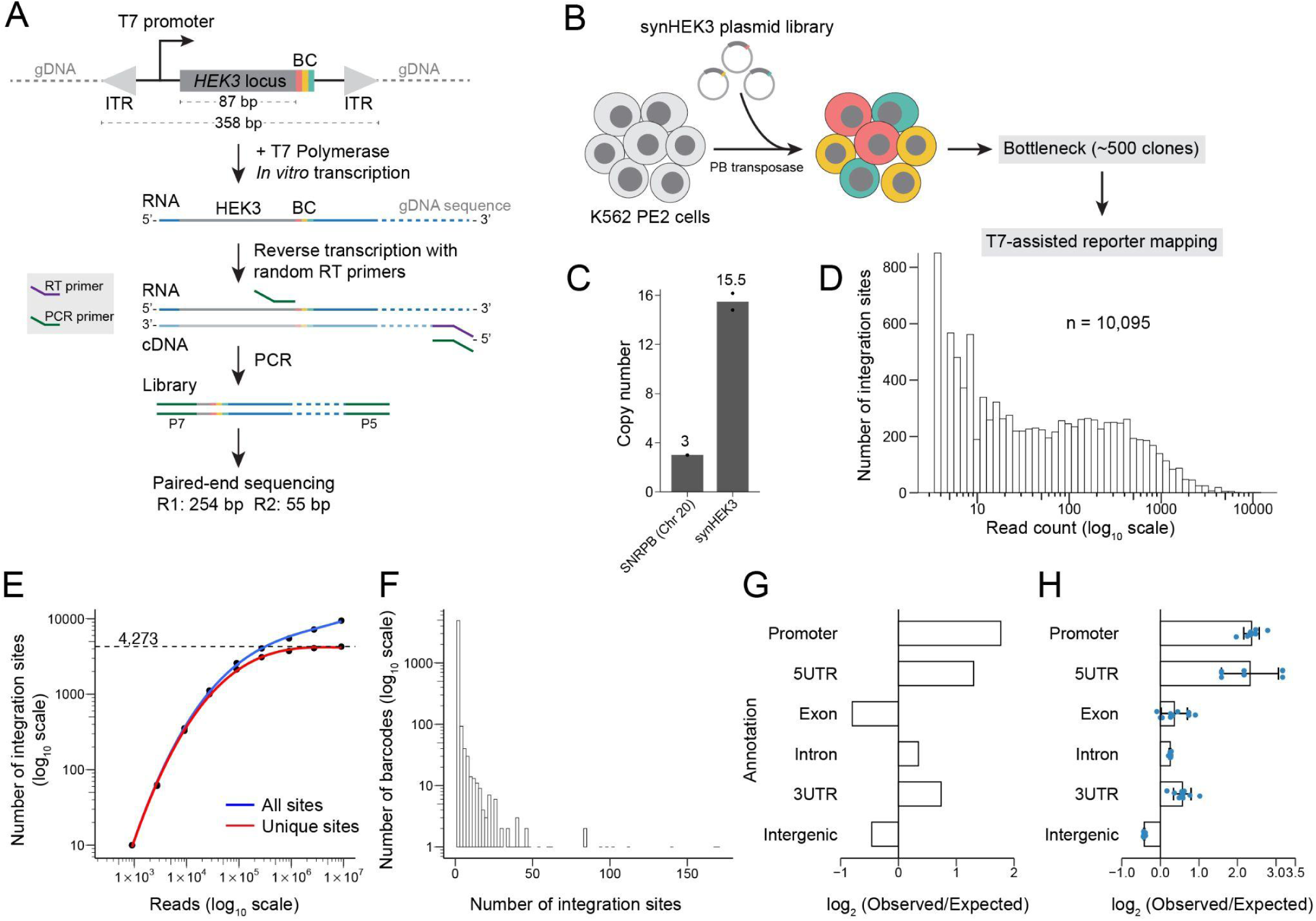
Characterization of synHEK3 insertion sites determined by T7-assisted reporter mapping assay. **A**) Schematic of the synHEK3 reporter construct and the T7-assisted reporter mapping method. **B**) Experimental workflow. A complex library of synHEK3 reporters was transfected into PE2-expressing K562 cells along with the *piggyBac* transposase. After transposons were stably integrated, the population of cells was bottlenecked to ∼500 clones. Cells derived from the expansion of this bottlenecked pool were then subjected to T7-assisted reporter mapping. **C**) Copy number of synHEK3 reporters was estimated by qPCR using that of *SNRPB* (3 copies) as a reference. **D**) Histogram of T7 mapping counts of all synHEK3 reporters (n = 10,095). *x*-axis on log_10_ scale. **E**) Scatter plot of number of integration sites recovered at different sequencing depths. Overlaid polynomial regression lines fit to all sites (blue) or unique (red) sites. *x*- and *y*-axis on log_10_ scale. **F**) Histogram of number of distinct genomic integration sites per synHEK3 barcode. *y*-axis on log_10_ scale. **G**) Log_2_ enrichment of synHEK3 reporter across major genomic features relative to expectation based on feature sizes. **H**) Log_2_ enrichment of synHEK3 reporter across major genomic features relative to expected number of TTAA motifs in each feature. Blue points: log_2_ enrichment calculated using 10 sets of randomly sampled TTAA motifs (n = 4,273). Error bar: standard deviation.

**Supplementary Figure 2.**
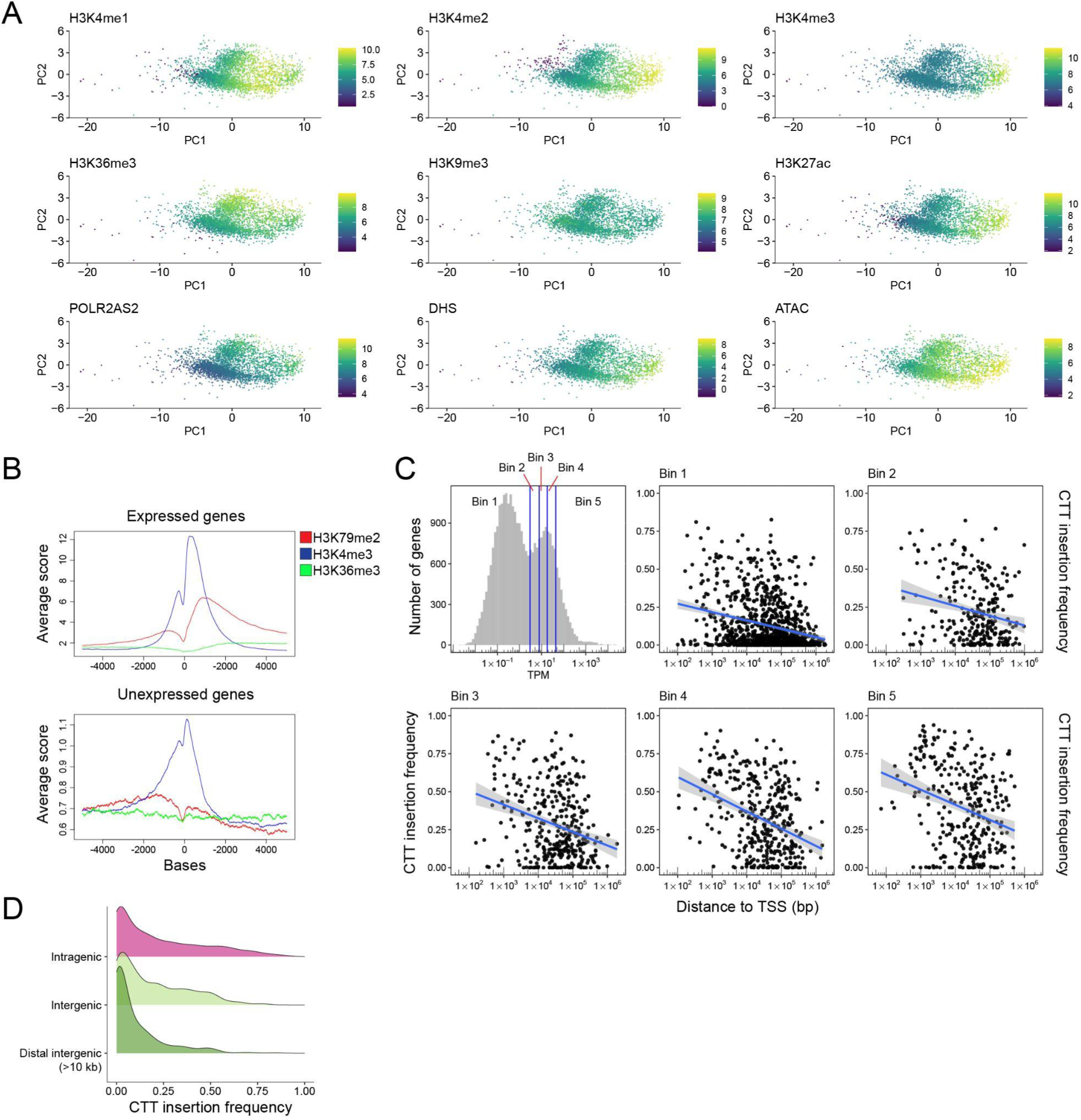
Chromatin context has a major impact on prime editing efficiency. **A**) PCA plots of synHEK3 reporters generated using chromatin scores as features. The first two PCs are plotted (PC1: variance 62%; PC2: variance 9%). Points are colored by the scores of indicated chromatin features. **B**) Histogram of ChIP-seq signal of H3K79me2 (red), H3K4me3 (blue) and H3K36me3 (green) in an 10-kb window surrounding the TSS of genes that are expressed (n = 11,514, TPM > 3), top) or unexpressed (n = 10,506, TPM < 3, bottom) in K562 cells. **C**) Prime editing efficiency for intragenic synHEK3 reporters as a function of their expression levels and distance to the TSS. The reporters were binned based on the TPM of genes underneath, as illustrated via the strategy in the top-left panel (histogram of TPM of all genes in K562 cells; *x*-axis: log_10_ scale). Bin 1 contains unexpressed genes, while Bin 2-5 are equal size bins of genes sorted by increasing expression levels. The remaining plots show the inverse correlation between distance to TSS (*x*-axis; log_10_ scale) and prime editing efficiency (y-scale). Blue line: linear regression line with confidence interval in gray. **D**) Density plot of CTT insertion frequency of synHEK3 reporter in intragenic, proximal intergenic and distal intergenic (>10 kb) regions.

**Supplementary Figure 3.**
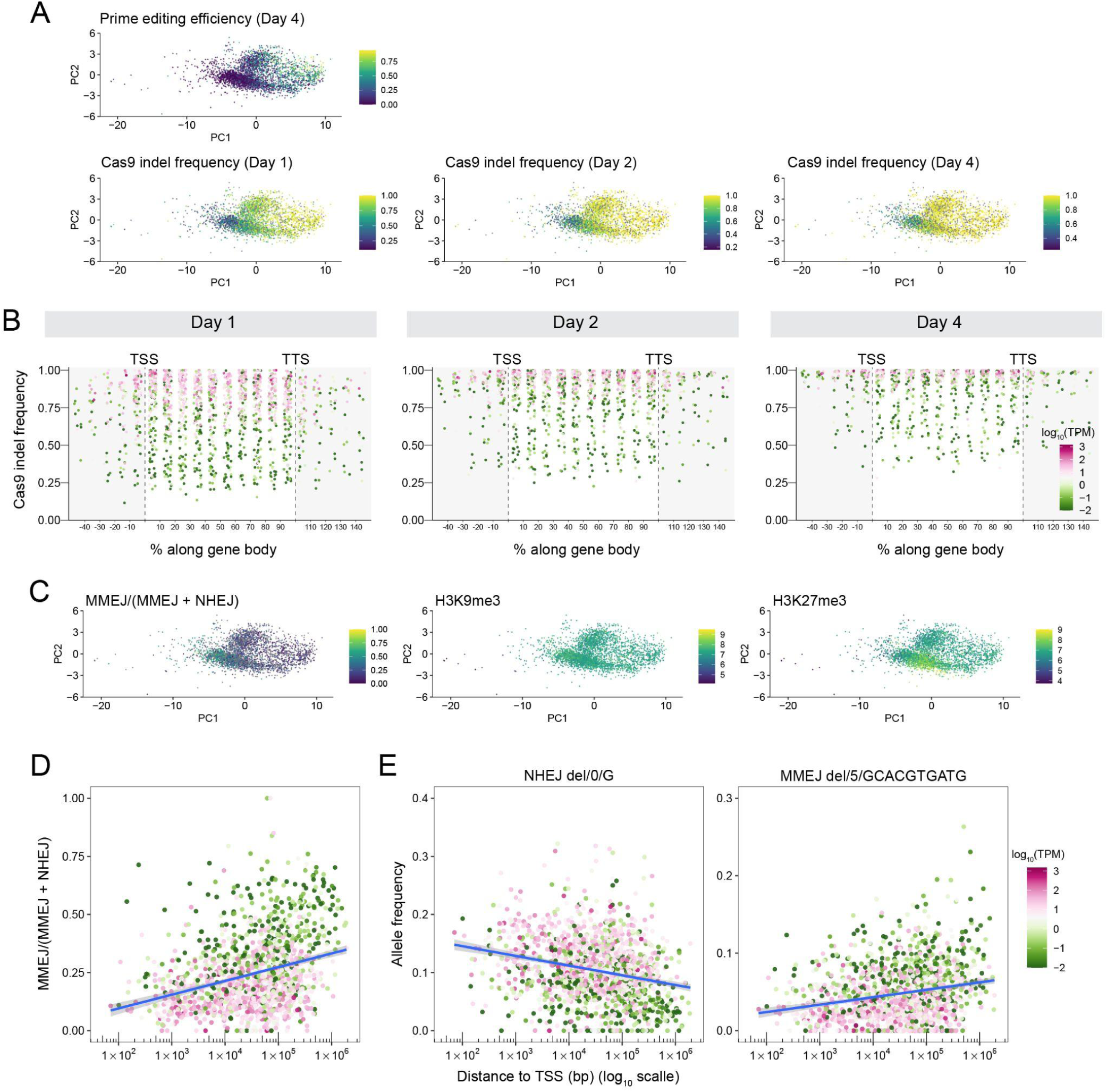
Comparison between prime editing and Cas9, leveraging a common set of integrated editing reporters. **A**) PCA plots of synHEK3 reporters generated using chromatin scores as features. The first two PCs are plotted (PC1: variance 62%; PC2: variance 9%). Points are colored by prime editing efficiency at Day 4 (top) and Cas9 indel frequency measured at Day 1, 2 and 4 (bottom). **B**) Cas9 indel frequency measured at Day 1, 2 and 4 for gene-proximal reporters. Distance was calculated relative to the closest TSS, scaled by gene length and binned. Points are colored based on expression levels (log_10_) of the genes. TPM, transcripts per million. **C**) PCA plots of synHEK3 reporters generated using chromatin scores as features. The first two PCs are plotted (PC1: variance 62%; PC2: variance 9%). Points are colored by MMEJ / (MMEJ + NHEJ) ratio and two heterochromatin markers (H3K9me3 and H3K27me3). **D-E**) Scatter plots of the MMEJ / (MMEJ + NHEJ) ratio or allele frequencies of intragenic synHEK3 reporters and their distance to corresponding TSSs (*x*-axis; log_10_ scale). Points are colored based on expression levels (log_10_) of the genes. TPM, transcripts per million. Blue line: linear regression line with confidence interval in gray. **D**) The MMEJ / (MMEJ + NHEJ) ratio is plotted. **E**) Allele frequencies of the most frequent NHEJ (left) and MMEJ (right) alleles are plotted. The “NHEJ del/0/G” allele contains a single-base deletion of G at the Cas9 cut site. The “MMEJ del/5/GCACGTGATG” allele contains a bidirectional deletion of a 10-bp sequence (GCACGTGATG) around the cutsite.

**Supplementary Figure 4.**
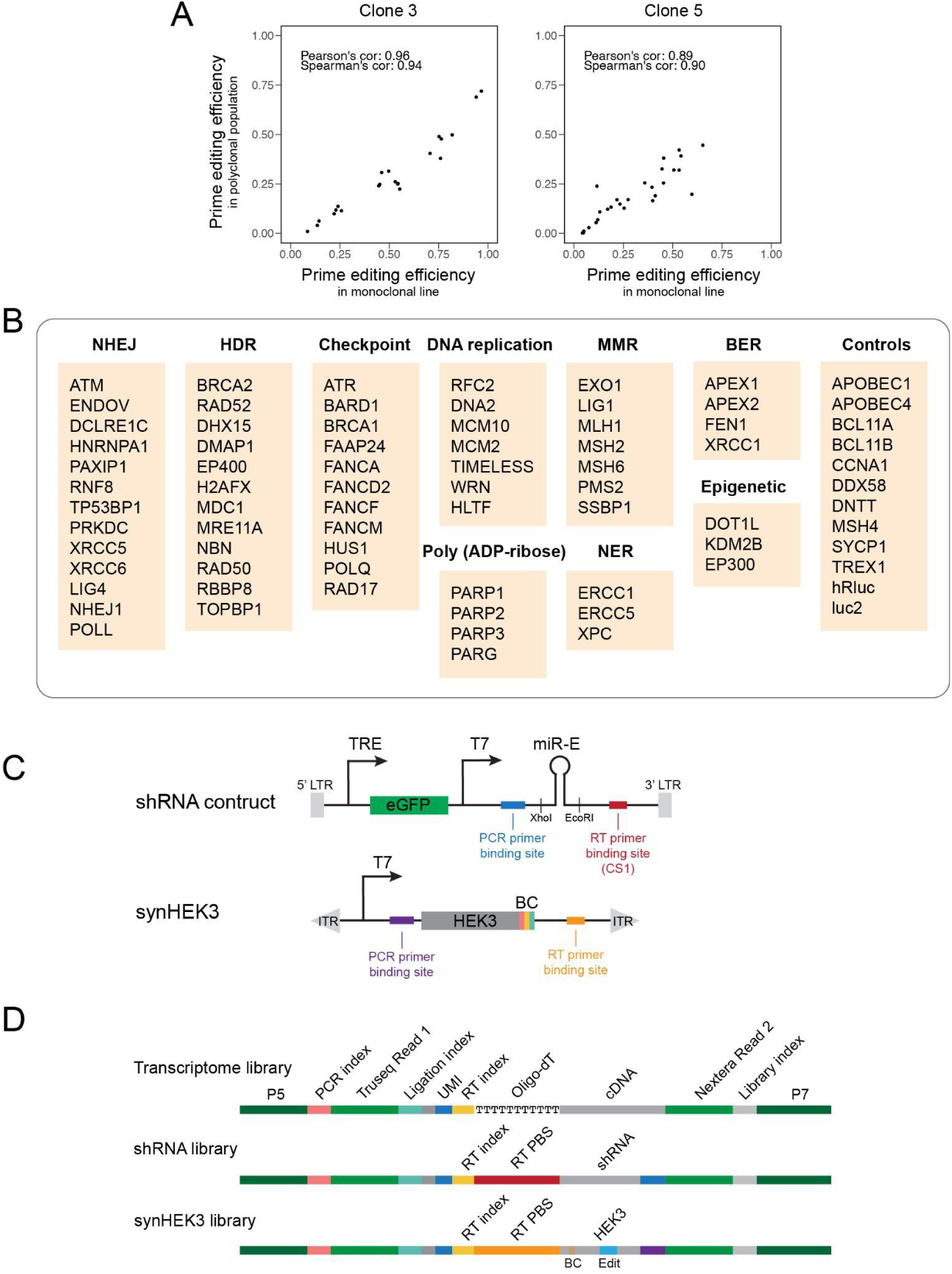
Experimental setup of the pooled shRNA screen and its readout via T7 IST-assisted sci-RNA-seq3. **A**) Correlation between efficiencies of random 3-bp insertions measured in the monoclonal lines versus efficiencies measured for the same reporters in the original polyclonal population. **B**) List of genes targeted by the shRNA library. Genes are grouped by pathways. Control genes are shown separately. **C**) Schematic of the lentiviral shRNA construct and the synHEK3 reporter, with key features relevant to the sci-RNA-seq3 workflow highlighted. TRE: tetracycline response element. **D**) Schematic structures of the sci-RNA-seq3, shRNA and synHEK3 libraries. UMI: unique molecular identifier; RT PBS: reverse transcription primer binding site.

**Supplementary Figure 5.**
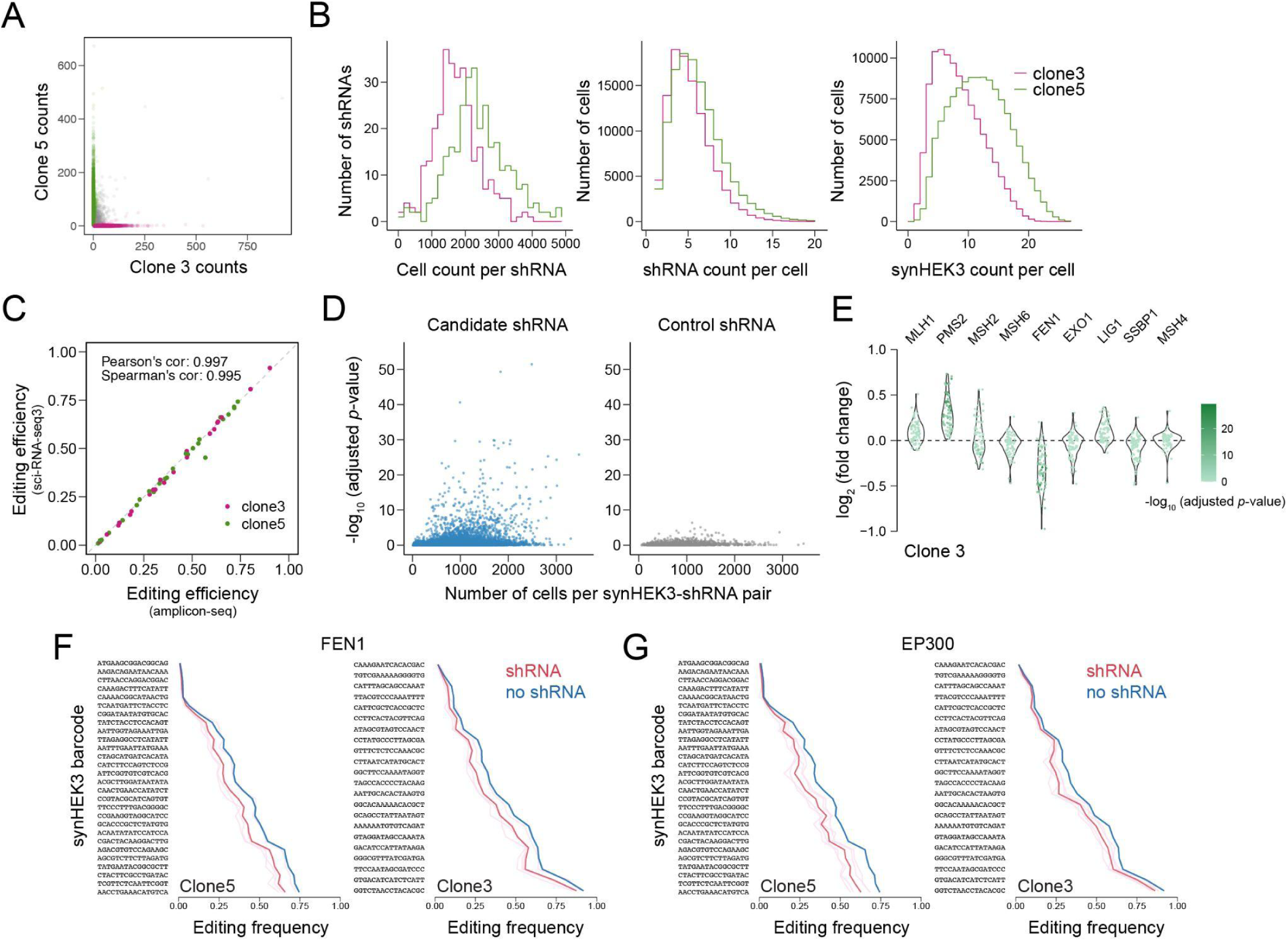
Pooled shRNA screen in sci-RNA-seq3. **A**) Scatter plot of synHEK3 UMIs detected in single cells in sci-RNA-seq3 data. Cells assigned to the two clones are colored (green: clone 5; pink: clone 3). Mixed cells are in gray. **B**) Histograms of cell count per shRNA (left), number of shRNAs captured per cell (middle), and number of synHEK3 reporters captured per cell (right) in the two clones. **C**) Scatter plot of prime editing frequencies of synHEK3 reporters estimated in sci-RNA-seq3 and in bulk amplicon sequencing. **D**) Scatter plots of the number of cells per synHEK3-shRNA pair and corresponding adjusted *p* values (-log_10_). Candidate shRNAs (left) and control shRNAs (right) are plotted separately. **E**) Effects of shRNAs targeting genes in the MMR pathway in clone 3. Log_2_ fold changes of prime editing efficiencies of synHEK3-shRNA pairs are plotted and colored by their corresponding adjusted *p* values (-log_10_). **F-G**) Effects of shRNAs against FEN1 and EP300. Pink lines: editing frequencies in cells with individual shRNAs; red line: mean editing frequencies of the gene-targeting shRNAs; light blue lines: control editing frequencies for individual shRNAs (not visible because low variance relative to mean line, shown in blue); blue line: mean control editing frequencies.

**Supplementary Figure 6.**
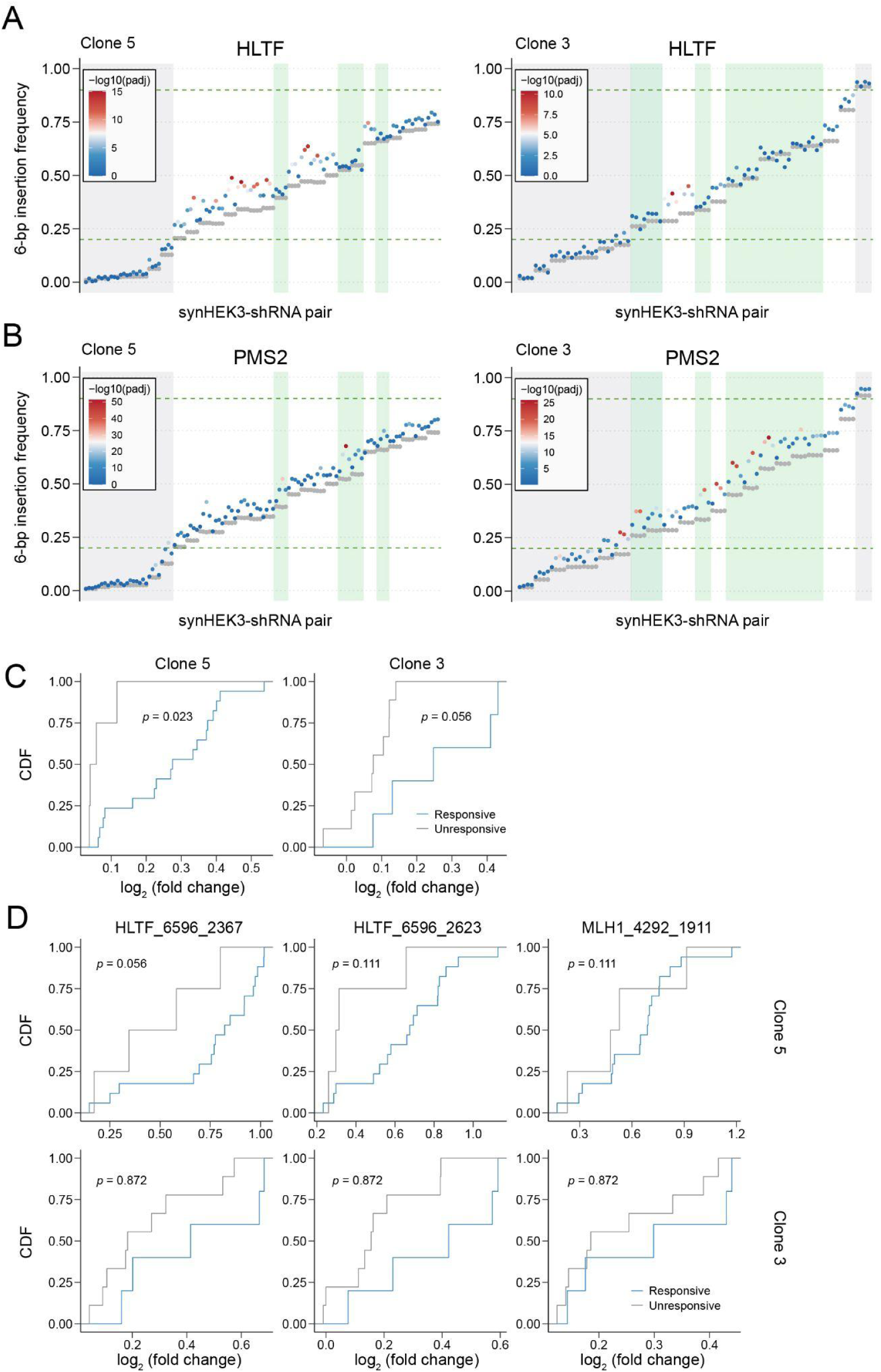
Chromatin context-specific response to HLTF inhibition. **A**) Effects of shRNAs against HLTF on all synHEK3 reporters in clone 5 (left) and clone 3 (right). For every synHEK3-shRNA pair, editing frequencies in cells with candidate shRNA are plotted and colored by their statistical significance. Control editing frequencies are shown in gray. SynHEK3 reporters that are less responsive to HLTF inhibition are highlighted in green. synHEK3 reporters with low (< 0.2) or very high (> 0.9) editing frequencies are colored in gray. Dashed horizontal lines indicate editing frequency of 0.2 and 0.9. **B**) Effects of shRNAs against PMS2 on all synHEK3 reporters in clone 5 (left) and clone 3 (right). For every synHEK3-shRNA pair, editing frequencies in cells with candidate shRNA are plotted and colored by their statistical significance. Control editing frequencies are shown in gray. SynHEK3 reporters that are less responsive to HLTF inhibition are highlighted in green. synHEK3 reporters with low (< 0.2) or very high (> 0.9) editing frequencies are colored in gray. Dashed horizontal lines indicate editing frequency of 0.2 and 0.9. **C**) Cumulative distribution function (CDF) plots of log_2_ fold change of mean editing frequency induced by shRNAs against HLTF. synHEK3 reporters are colored by their responsiveness to HLTF inhibition based on the shRNA screen. *P* value: two-sided Kolmogorov–Smirnov test. **D**) Validation of differential responsiveness to HLTF inhibition. CDF plots of log_2_ fold change of editing frequency induced by shRNAs against HLTF (HLTF_6596_2367: used in the shRNA screen; HLTF_6596_2623: an orthogonal shRNA) or MLH1 (MLH1_4292_1911). synHEK3 reporters are colored by their responsiveness to HLTF inhibition based on the shRNA screen. *P* value: two-sided Kolmogorov–Smirnov test.

**Supplementary Figure 7.**
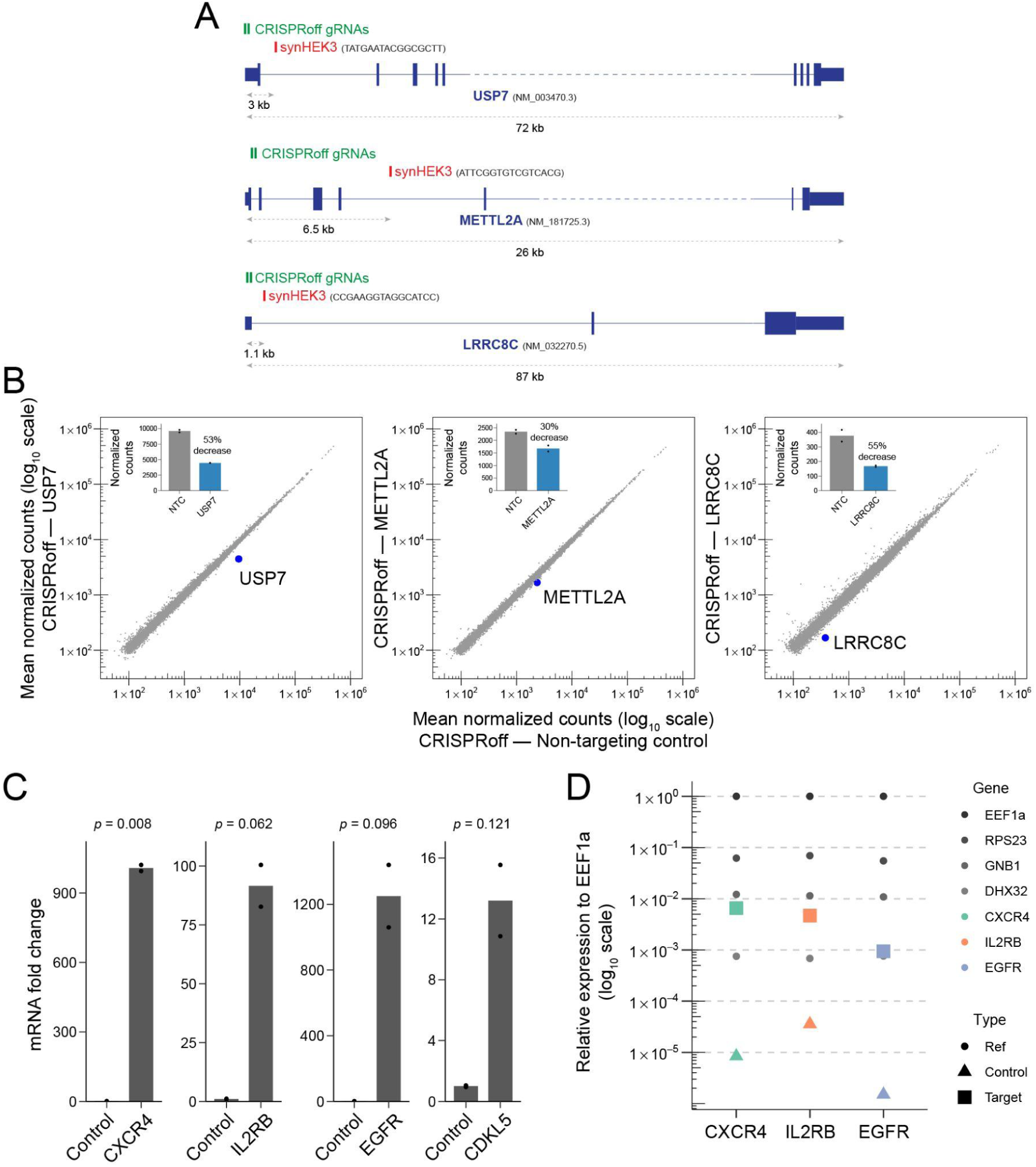
Modulating prime editing outcomes by epigenetic reprogramming. **A**) Schematic diagrams of target genes in the CRISPRoff experiment, with the locations of the CRISPRoff gRNAs and synHEK3 reporters annotated. **B**) Scatter plots of mean normalized counts of gene expression in cells transfected with CRISPRoff gRNAs at Day 11 (2 replicates each). *x*- and *y*-axis on log_10_ scale. Insets are bar plots of normalized counts of the CRISPRoff target genes. NTC: non-targeting control. **C**) mRNA fold-change of CRIPSRa target genes quantified by qPCR. *P* value: welch’s two sample t-test. **D**) Relative expression levels of CRISPRa target genes compared to a set of reference genes. Circle: reference genes; triangle: endogenous expression levels of the target genes; square: expression levels of the target genes after CRISPR activation. *y*-axis on log_10_ scale.

