## Supplementary tables for "Chromatin context-dependent regulation and epigenetic manipulation of prime editing": KEY RESOURCES TABLE.docx

| REAGENT or RESOURCE | SOURCE | IDENTIFIER |
| --- | --- | --- |
| Critical Commercial Assays | | |
| TruSeq RNA Library Prep Kit v2 | Illumina | Cat#RS-122-2001 |
| TruSeq Stranded mRNA kit | Illumina | Cat#20020594 |
| Illumina TruSeq RNA UD Indexes | Illumina | Cat#20023785 |
| NextSeq 1000/2000 P2 Reagents (100 Cycles) v3 | Illumina | Cat#20046811 |
| NextSeq 500/550 Mid Output Kit v2.5 (150 Cycles) | Illumina | Cat#20024904 |
| MiSeq Reagent Kit v2 (300-cycles) | Illumina | Cat#MS-102-2002 |
| MiSeq Reagent Kit v3 (150-cycle) | Illumina | Cat#MS-102-3001 |
| MiSeq Reagent Kit v2 (50-cycles) | Illumina | Cat#MS-102-2001 |
| Bacterial and Virus Strains | | |
| NEB Stable Competent *E. coli* | New England Biolabs | Cat#C3040H |
| NEB 10-beta Electrocompetent *E. coli* | New England Biolabs | Cat#C3020K |
| Chemicals, peptides, and recombinant proteins | | |
| RPMI 1640 Medium | Gibco | Cat#11875119 |
| DMEM, High Glucose | Gibco | Cat#11965118 |
| FBS | Hyclone | Cat#SH30396.03 |
| Penicillin-Streptomycin (10,000 U/mL) | Gibco | Cat#15140122 |
| DNeasy Blood & Tissue Kit | QIAGEN | Cat#69504 |
| HiScribe™ T7 High Yield RNA Synthesis Kit | New England Biolabs | Cat#E2040S |
| HiScribe^®^ T7 Quick High Yield RNA Synthesis Kit | New England Biolabs | Cat#E2050 |
| RNase-Free DNase Set | QIAGEN | Cat#79254 |
| TURBO™ DNase | Invitrogen | Cat#AM2238 |
| TRIzol LS Reagent | Invitrogen | Cat#10296010 |
| RNeasy Mini Kit | QIAGEN | Cat#74104 |
| Glycogen (5 mg/mL) | Invitrogen | Cat#AM9510 |
| SuperScript™ IV Reverse Transcriptase | Invitrogen | Cat#18091050 |
| RNaseOUT™ Recombinant Ribonuclease Inhibitor | Invitrogen | Cat#10777019 |
| Proteinase K | Thermo Scientific | Cat#EO0491 |
| AMPure XP Reagent | Beckman Coulter Life Sciences | Cat#A63882 |
| KAPA2G Robust HotStart ReadyMix PCR Kit | Roche | Cat#KK5702 |
| Power SYBR™ Green PCR Master Mix | Applied Biosystems | Cat#4367659 |
| NEBuilder^®^ HiFi DNA Assembly Cloning Kit | New England Biolabs | Cat#E5520S |
| BsmBI-v2 | New England Biolabs | Cat#R0739S |
| FastDigest BpiI (IIs class) | Thermo Scientific | Cat#FD1014 |
| I-SceI | New England Biolabs | Cat#R0694S |
| XhoI | New England Biolabs | Cat#R0146S |
| EcoRI-HF | New England Biolabs | Cat#R3101S |
| BamHI-HF | New England Biolabs | Cat#R3136S |
| XbaI | New England Biolabs | Cat#R0145S |
| SpeI-HF | New England Biolabs | Cat#R3133S |
| NotI-HF | New England Biolabs | Cat#R3189S |
| T4 DNA ligase | New England Biolabs | Cat#M0202L |
| SF Cell Line 4D-Nucleofector™ X Kit | Lonza Bioscience | Cat#V4XC-2012 |
| SF Cell Line 96-well Nucleofector™^M^ Kit | Lonza Bioscience | Cat#V4SC-2096 |
| ViraPower™ Lentiviral Packaging Mix | Invitrogen | Cat#K497500 |
| Lipofectamine™ 3000 Transfection Reagent | Invitrogen | Cat#L3000001 |
| PEG-it™ Virus Precipitation Solution (5×) | System Biosciences | Cat#LV810A-1 |
| Polybrene | Millipore | Cat#TR-1003-G |
| Geneticin | Gibco | Cat#10131035 |
| Blasticidin S HCl (10 mg/mL) | Gibco | Cat#A1113903 |
| DEPC (Diethyl Pyrocarbonate) | Millipore Sigma | CAt#D5758-25ML |
| YOYO™-1 Iodide (491/509) - 1 mM Solution in DMSO | Invitrogen | Cat#Y3601 |
| dNTP mix | New England Biolabs | Cat#N0447L |
| SuperScript™ IV Reverse Transcriptase | Invitrogen | Cat#18090050 |
| NEBNext^®^ Ultra™ II Non-Directional RNA Second Strand Synthesis Module | New England Biolabs | Cat#E6111L |
| Protease | QIAGEN | Cat#19157 |
| Buffer EB | QIAGEN | Cat319086 |
| Tn5 transposase | Diagenode | Cat#C01070010-20 |
| BSA | New England Biolabs | Cat#B9000S |
| Monarch® DNA Gel Extraction Kit | New England Biolabs | Cat#T1020S |
| NEBNext^®^ High-Fidelity 2X PCR Master Mix | New England Biolabs | Cat#M0541L |
| OneTaq® Hot Start 2X Master Mix with Standard Buffer | New England Biolabs | Cat#M0484L |
| SYBR™ Green I Nucleic Acid Gel Stain | Invitrogen | Cat#S7563 |
| Alt-R™ S.p. Cas9 Nuclease V3 | Integrated DNA Technologies | Cat#1081059 |
| Alt-R® Cas9 Electroporation Enhancer, 2 nmol | Integrated DNA Technologies | Cat#1075915 |
| Deposited data | | |
| GSE228465 |  |  |
| Experimental models: Cell lines | | |
| K562 PE2-Puro | Choi et al., 2021 | NA |
| K562 dCas9-VP64 | Chardon et al., 2023 | NA |
| Oligonucleotides | | |
| Table S1 | NA | NA |
| Recombinant DNA | | |
| PB-T7-HEK3-BC | This paper | NA |
| LT3-GFP-T7-miR-E-CS1-PGK-Neomycin | This paper | NA |
| Lenti-rtTA-P2A-Blast | This paper | NA |
| pU6-Sp-pegRNA-HEK3-ins6N | This paper | NA |
| pU6-Sp-dual-gRNA | This paper | NA |
| pU6-Sp-gRNA-2XMS2 | This paper | NA |
| PB-CMV-MCP-XTEN80-p65-Rta-3xNLS-P2A-T2A-mPlum | This paper | NA |
| pU6-Sp-pegRNA-HEK3-ins3N | This paper | NA |
| pU6-Sp-pegRNA-HEK3-insCTT | Anzalone et al., 2019 | Addgene: 132778 |
| pCMV-PEmax | Chen et al., 2021 | Addgene: 174820 |
| CRISPRoff-v2.1 | Nuñez et al., 2021 | Addgene: 167981 |
| Software and algorithms | | |
| Bcl2fastq (v2.20) | Illumina | https://support.illumina.com/sequencing/sequencing_software/bcl2fastq-conversion-software.html |
| Samtools (v1.9) | Li et al., 2009 | http://www.htslib.org |
| Bedops (v2.4.35) | Neph et al., 2012 | https://bedops.readthedocs.io/en/latest/index.html |
| needleall | Needleman et al., 1970  Kruskal, 1983 | https://emboss.sourceforge.net/apps/release/6.6/emboss/apps/needleall.html |
| STAR (v2.7.6a) | Dobin et al., 2013 | https://github.com/alexdobin/STAR |
| Salmon (v1.9.0) | Patro et al., 2017 | https://salmon.readthedocs.io/en/latest/ |
| Cutadapt (v4.1) | Martin, 2011 | https://cutadapt.readthedocs.io/en/stable/ |
| TrimGalore (v0.6.6) | Krueger, 2012 | https://www.bioinformatics.babraham.ac.uk/projects/trim_galore/ |
| Bwa (v0.7.17) | Li et al., 2009 | https://bio-bwa.sourceforge.net/ |
| GenomicRanges | Lawrence et al., 2013 | https://bioconductor.org/packages/release/bioc/html/GenomicRanges.html |
| ChIPseeker | Yu et al., 2015 | https://bioconductor.org/packages/release/bioc/html/ChIPseeker.html |
| Homer | Heinz et al., 2010 | http://homer.ucsd.edu/homer/ |
| Genomation | Akalin et al., 2015 | https://bioconductor.org/packages/release/bioc/html/genomation.html |
| HTseq (v.2.0.2) | Putri et al., 2022 | https://htseq.readthedocs.io/en/master/ |
| Seurat (v4.0.0) | Hao et al., 2021 | https://satijalab.org/seurat/ |
| Betareg (v3.1.4) | Cribari-Neto et al., 2010 | https://cran.r-project.org/web/packages/betareg/index.html |
| DESeq2 | Love et al., 2014 | https://bioconductor.org/packages/release/bioc/html/DESeq2.html |
| WebLogo 3 | Crooks et al., 2004 | https://weblogo.threeplusone.com/ |
| ComplexHeatmap | Gu et al., 2016 | https://bioconductor.org/packages/release/bioc/html/ComplexHeatmap.html |
| IGV | Robinson et al., 2011 | http://software.broadinstitute.org/software/igv/ |
